# Structural Basis of Nidovirus Replication Organelle Evolution Revealed by the Arterivirus DMV Pore Complex

**DOI:** 10.64898/2026.06.15.732489

**Authors:** Wenxin Zhang, Tingting Yang, Wanlong Hu, Le Zheng, Yixin Huang, Lijie Zhong, Qiannan Li, Yuxin Gao, Qian Yang, Yanfei Wang, Haibo Jiang, Xiulian Yu, Tao Ni

## Abstract

Positive-strand RNA viruses of the order Nidovirales – which include coronaviruses and arteriviruses – remodel host membranes into double-membrane vesicles (DMVs) as replication organelle to shield viral RNA synthesis from immune sensors. In coronaviruses, newly synthesized RNA is exported to the cytoplasm through a massive DMV pore complex formed by viral non-structural proteins (nsps). However, it remains unknown whether this elaborate architecture is unique to large-genome coronaviruses or a universal hallmark of the order. Here, we integrate *in situ* cryo-electron tomography and single-particle cryo-electron microscopy to resolve the atomic structure of DMV pore complex within its native membrane environment, from Equine Arteritis Virus (EAV), a prototype arterivirus with small genome. Despite lacking obvious sequence homology, the minimal EAV pore shares conserved architectural principles with its elaborate coronavirus counterpart. EAV pore complex is formed by a 12:12 stoichiometry of nsp2 and nsp3 protomers organized into four stacked concentric rings on double-membrane, yet generating pronounced structure symmetry mismatch. Functionally, the complex displays a distinct pore profile while preserving a positively charged central channel, essential for viral replication and transport. These findings demonstrate that, despite diversity in genome size and virion morphology, nidovirus replication organelles exhibit striking evolutionary conservation at the atomic structural level. Collectively, we propose that the order Nidovirales can be unified at the ultrastructural level by this conserved signature pore complex on the DMV-based replication organelle.

## BACKGROUND

Positive-strand RNA viruses commonly remodel host intracellular membranes to form specialized replication organelles (ROs), which concentrate viral and host factors to facilitate efficient viral RNA synthesis while physically shielding viral components from cytoplasmic innate immune sensors^1–7^. This strategy is employed across diverse virus families, including coronaviruses, flaviviruses, and alphaviruses^8–10^. Among the most structurally complex ROs are the double-membrane vesicles (DMVs) induced by members of the *Nidovirales* order^11^, which include the clinically significant viruses in *Coronaviridae* (e.g., SARS-CoV-2, MERS-CoV), the economically important *Arteriviridae* (e.g., Equine arteritis virus (EAV), Simian hemorrhagic fever virus (SHFV), and Porcine reproductive and respiratory syndrome virus (PRRSV)), as well as other families such as *Roniviridae* and *Mesoniviridae*^12^*, etc*. Despite marked differences in genome size, host range, and pathogenicity, nidoviruses share a conserved replication mechanism centered on the formation of DMVs that house the viral replication-transcription complexes^12,13^.

A central, unresolved question in nidovirus replication is how newly synthesized, positive-sense genomic and subgenomic RNAs are exported from the seemingly sealed lumen of DMVs to the cytoplasm for translation and virion assembly. In coronaviruses, previous *in situ* cryo-electron tomography (cryo-ET) studies of mouse hepatitis virus (MHV) and SARS-CoV-2-infected cells first visualized a putative RNA export channel within their replication organelles^8,14^. These studies confirmed the existence of a pore-like complex, exhibiting six-fold symmetry and spanning the double membranes of the DMVs. The atomic molecular architecture of the SARS-CoV-2 nsp3–nsp4 pore complex was then resolved by cryo-ET and subtomogram averaging (STA) in isolated recombinant DMVs^15^. This structure provided the atomic-level description for DMV pore complex in coronaviruses, revealing a unique stoichiometry – 12 nsp3 and 12 nsp4 forming 4-layer of longitudinal but interdigitated hexamer rings. Functionally, the complex is a putative RNA translocation pore coordinated by a central positively charged arginine ring, essential to virus replication.

The complexity and the large size of the coronavirus DMV pore complex raise an important question. Is this architectural solution unique to coronaviruses or shared by more divergent members of the *Nidovirales* order? How is the functional architecture of this essential RNA conduit maintained across the entire viral order despite extreme sequence divergence? Arteriviruses are particularly informative in this context because they possess much smaller genomes than coronaviruses, approximately 10–15 kb compared with ∼30 kb^13,16–19^. Thus, comparing arteriviral and coronaviral replication organelles provides an opportunity to assess whether a related pore architecture is conserved across deeply divergent nidovirus lineages. Furthermore, Arteriviruses are also biologically and economically important^20–23^. PRRSV causes devastating disease in swine worldwide^24–26^. SHFV can cause severe, often fatal disease in non-human primates, and has been used as a model for viral hemorrhagic fever^23,27^. EAV, the prototype of the family, is the etiologic agent of equine viral arteritis^28,29^. EAV serves as a simplified model for nidovirus study due to its smallest genome (approximately 12.7 kb) among nidoviruses, yet it induces biochemically analogous ER-derived DMVs for replication^30–32^. Its transmembrane replicase subunits nsp2 and nsp3 are necessary and sufficient to drive DMV biogenesis, paralleling the role of coronavirus nsp3 and nsp4 despite the absence of obvious sequence homology^11,33–36^. Recent cryo-ET studies of arterivirus-infected cells provided initial evidence for the existence of structurally similar DMV pores, suggesting that DMV pores may be conserved beyond coronaviruses^37^. However, whether these arteriviral pores share a related molecular architecture with coronavirus DMV pores remains unclear.

Here, we characterize EAV replication organelle *in situ* by cryo-ET and report structure of the EAV DMV pore complex by cryo-electron microscopy (cryo-EM). This work provides the first atomic resolution view of an arterivirus replicopore, revealing the minimal essential machinery that serves as a prototype model for nidoviruses. Through detailed structural analysis, replicon-based functional assays, and comparison with the SARS-CoV-2 DMV pore complex, we elucidate both divergent and conserved principles of membrane remodeling and RNA transport between two highly divergent nidovirus lineages. While we observe substantial differences in the overall pore profile and complex architecture, the core molecular principles – including subunit stoichiometry, protein complex topology, and a charged central channel compatible with RNA transport – remain strictly conserved across these two distant viral families. We therefore propose that the nidovirus replication organelle is fundamentally defined as a broadly conserved, DMV-based compartment characterized by this signature pore complex. Our findings not only advance the mechanistic understanding of replication organelle biogenesis but also identify conserved architectural features that could serve as foundational targets for broad-spectrum antiviral strategies.

## RESULTS

### Cryo-ET analysis of DMV pore complexes in EAV-infected cells

Nidoviruses are defined primarily by their genome organization and replication strategy^12,38,39^. They use a unique RNA replication and transcription mechanism to produce a nested set of 3′-coterminal subgenomic mRNAs during replication. In addition, they encode a large replicase polyprotein from ORF1a/ORF1b, regulated via a −1 ribosomal frameshift during translation. These replicase polyproteins, encoding a series of non-structural proteins (nsps), mediates virus genome replication within a specific DMV-based replication organelle.

To explore the replication organelle formation across the *Nidovirales* order, we first characterized EAV virus replication and DMV formation *in situ* by cryo-focused ion beam milling and cryo-ET 12 hours post infection (**Extended Data Fig. 1a,b; Table 1**). Our results reveal marked difference compared with coronavirus (CoV) infection^14^: EAV DMVs are variable in size and morphology **(Fig. 1a-c)**, and DMVs with two inner vesicles sharing the same outer membrane are commonly observed (**Fig. 1a-c**). Notably, DMV pore complexes were observed spanning the junction of the two membranes (**Fig. 1c-d**). Tomographic segmentation highlighted pore complex on these DMVs, as well as the RNA inside the DMVs. Rod-shaped RNP-like structures were frequently present around the vesicles, similar to the RNP densities in virus particles (**Fig. 1a,b**; **Extended Data Fig. 1c,d**).To resolve DMV pore architecture during virus replication *in situ*, we performed subtomogram averaging (STA) and obtained ∼2 nm resolution density with C6 symmetry (**Fig. 1d,e; Extended Data Fig. 1e-f**). The averaged map revealed a compact, ring-like assembly centered on a membrane-spanning channel. The complex bridged the double-membrane interface with only a limited protrusion above the membrane (**Fig. 1d,e**), and was significantly shorter than its CoV counterpart^15,33^. The limited resolution and almost no sequence similarity among arterivirus and CoV DMV pore components preclude the confident modelling of such complicated structures. These structural features indicate that, despite substantial divergence in size and overall dimensions, EAV DMVs possess a conserved membrane-spanning pore architecture reminiscent of that observed in CoV.

**Fig. 1.**
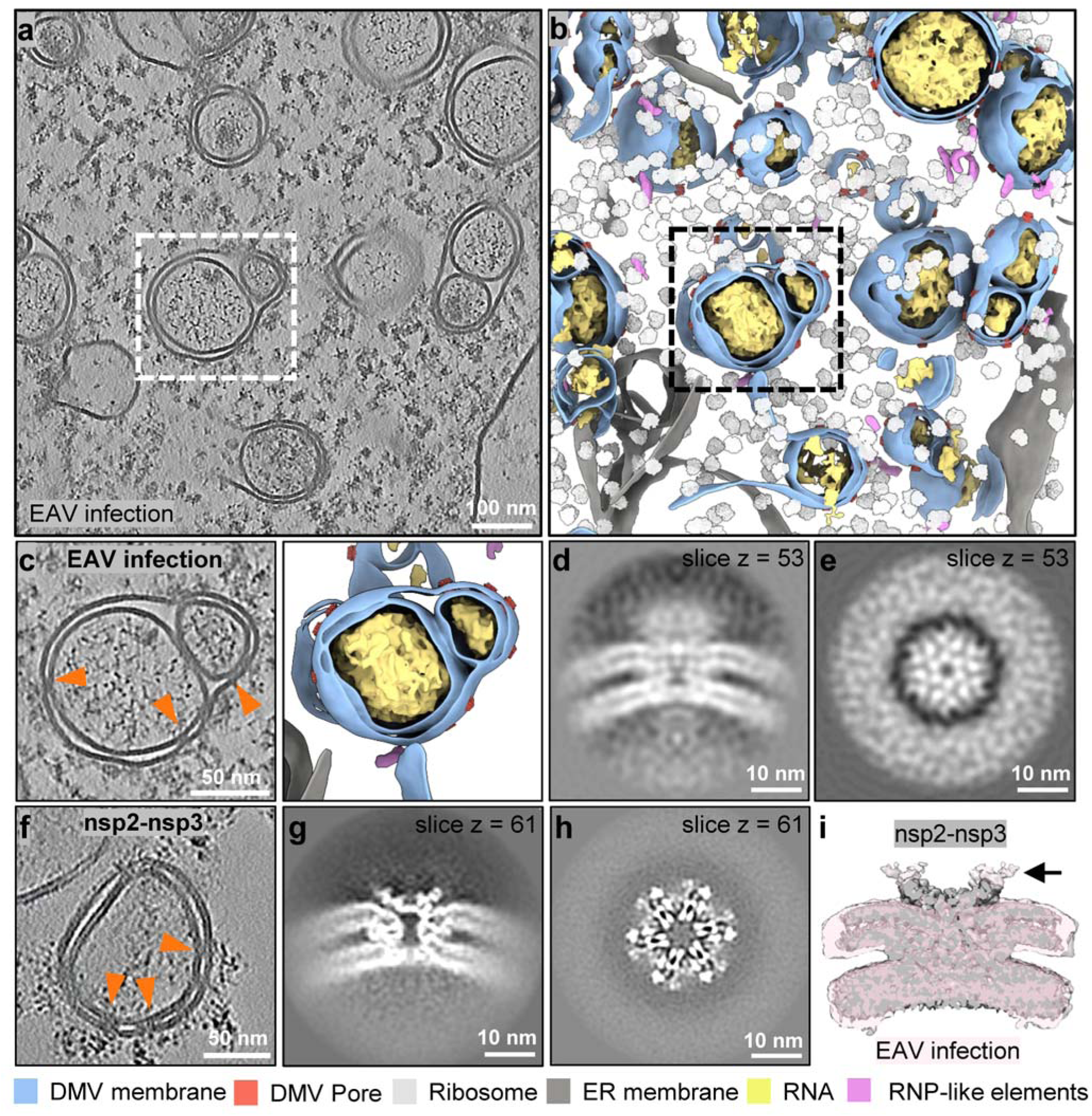
| Cryo-ET analysis of EAV-induced double-membrane vesicles and their pore complexes. **a**, Representative cryo-electron tomographic slice of an EAV-infected cell showing multiple double-membrane vesicles (DMVs). Dashed boxes indicate regions enlarged in c (left). **b,** Segmented rendering of the tomogram shown in a, with DMV membranes in blue, surrounding ER membranes in dark grey, ribosomes in light grey, luminal RNA densities in yellow, pore complexes in orange, and rod-shaped RNP-like elements in pink. Dashed boxes indicate regions enlarged in c (right). **c,** Enlarged views of selected DMVs and the corresponding segmentations, highlighting DMV morphology and DMV pore complexes (orange arrowheads). **d,e,** Subtomogram average of the EAV DMV pore complex from infected cells, shown in side view **(d)** and top view **(e). f,** Cryo-electron tomographic slice of a DMV induced by expression of EAV nsp2–nsp3 in HEK293F cells, with pore complexes indicated by orange arrowheads. **g, h,** Subtomogram average of the recombinant EAV DMV pore complex, shown in side view **(g)** and top view **(h). i,** Overlay of the subtomogram averages from recombinant DMVs (grey) and EAV infection-derived DMVs (pink), showing close agreement in overall architecture and dimensions. The arrows indicate the extra densities of pore complex in EAV infection.

To determine the high-resolution structure of EAV pore complex, we engineered a Twin-Strep-nsp2-3-FLAG construct to express and purify recombinant DMVs from EAV, PRRSV and SHFV utilizing a strategy following our previous work on CoV DMVs^15^. While these viruses exhibit distinct host preferences, their DMVs and pore complexes can be generated in HEK 293F cells (**Extended Data Fig. 2a-c**). We proceeded with the EAV construct for high-resolution structure determination. Since the EAV DMV pore complex is embedded into two membranes, it is not readily distinguishable in cryo-EM projection images, we initially employed cryo-ET and STA as previously. Iterative alignment and refinement yielded a resolution of 7.5 Å (**Fig. 1f-h; Extended Data Fig. 2d-j; Table 1**). We superimposed the two density maps of pore complexes from EAV infection-derived DMVs and recombinant DMVs, which revealed high overall similarity (**Fig. 1i**), although the EAV infection-derived map displayed an extra density extending from the cytoplasmic side of DMV membrane. These results indicate that the recombinant DMVs reproduce the core architecture of the *in situ* EAV pore complex and provide a suitable platform for subsequent high-resolution structural analysis.

### Structure of the EAV DMV pore complex in native membrane environment

Although the 7.5 Å resolution map reveals secondary structural elements, such as transmembrane helices, and accommodates the fitting of nsp3-Y1 hexamers, the lower half of the pore complex remained poorly resolved. Furthermore, unambiguous model assignment was not possible, as the structures of these components – either individually or as a complex – could not be reasonably predicted by AlphaFold^40^. To overcome the resolution limitations of cryo-ET STA and achieve near-atomic resolution, we transitioned to cryo-EM single-particle analysis (SPA). Using the 7.5 Å resolution STA density map as the template for high-resolution template matching, we revealed clear cross-correlation peaks along the DMV membranes in the purified samples (**Fig. 2b-c, Extended Data Fig. 3a & 4a**). Following two rounds of template matching, extensive classification and refinement with C6 symmetry, we achieved an initial overall resolution of 4 Å (**Extended Data Fig. 3a-g**). Subsequent relaxation of symmetry revealed that the basal portion of the density map adopts a C3 symmetry, analogous to the previously resolved CoV DMV pore complex. By aligning and merging the C3-symmetrized density maps in two opposite orientations (180° apart), we obtained a interpretable map at an overall resolution of 3 Å, with local resolutions ranging from 2.5 to 4.5 Å (**Fig. 2d-g, Extended Data Fig. 3h-j and Extended Data Fig. 4b,d**). A highly curved membrane junction fuses the inner and outer DMV membranes to form the pore complex (**Fig. 2d**).

**Fig. 2.**
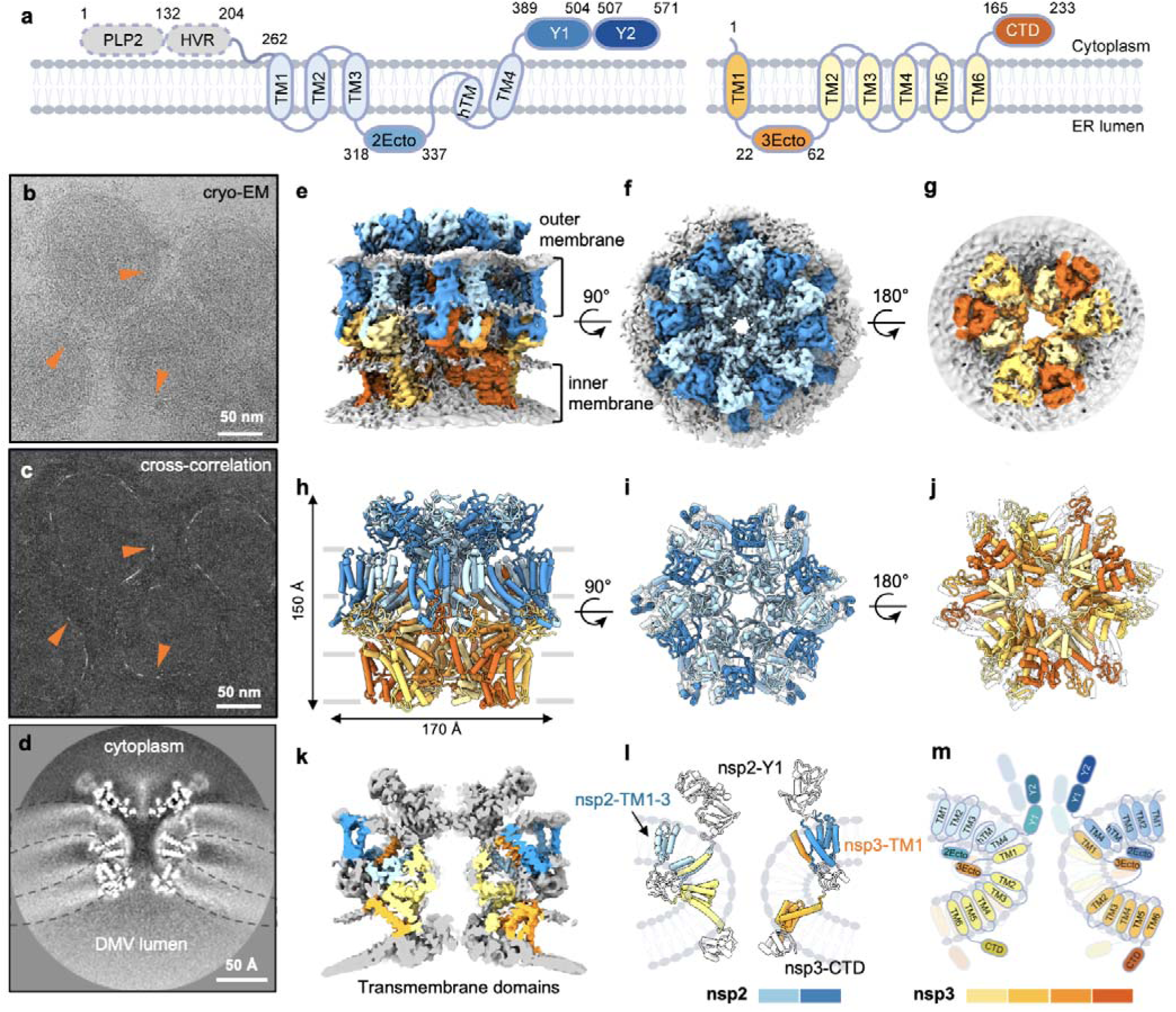
| Single-particle cryo-EM analysis of the EAV DMV pore complex. **a**, Schematic topology and domain organization of EAV nsp2 and nsp3 in the membrane bilayer, with amino acid numbers at domain boundaries indicated. The unresolved nsp2 PLP2 and hypervariable region (HVR) are shown as grey dashed ovals. TM, Transmembrane helices; 2Ecto, nsp2 ectodomain; 3Ecto, nsp3 ectodomain; CTD, C-terminal domain. **b,** A representative micrograph of purified EAV DMVs with the pore complexes indicated by orange arrowheads. **c,** Cross-correlation map identified by template-matching in *cis*TEM. Orange arrowheads indicate peak positions of detected DMV pore complexes. **d,** Longitudinal section through the cryo-EM density map of EAV DMV pore complex, with the curved double membranes indicated by dashed lines. **e–g,** Cryo-EM density map of the EAV DMV pore complex in side (**e**), cytoplasmic/top (**f**) and DMV luminal/bottom (**g**) views. Nsp2 protomer densities are coloured in alternating cyan and blue, nsp3 protomer densities in alternating yellow and orange, and membrane density in grey. **h–j,** Atomic model of the DMV pore complex fitted into the density map. The complex contains 12 copies of nsp2 and 12 copies of nsp3. **k,** Longitudinal cross-section of the density map showing the central channel of the pore complex traversing the paired DMV membranes, with transmembrane helices coloured. **l,** Side view of two pairs of nsp2–nsp3 heterprotomers within the double-membrane context highlighting the arrangement of transmembrane helices. Left, inner protomer (nsp2I and nsp3I); Right, outer protomer (nsp2O and nsp3O). **m,** A schematic model showing the topology and domain organization of nsp2 and nsp3 within the EAV DMV pore complex. The nsp2 inner protomer (nsp2I, left) is cyan and outer protomer (nsp2O, right) is blue; the nsp3 inner protomer (nsp3I, left) is yellow and outer protomer (nsp3O, right) is orange. Scale bars are indicated in the micrographs.

We next built an atomic model *de novo*, utilizing AlphaFold-predicted domains for cross reference^40^. All transmembrane helices and domains were clearly resolved, with the exception of the N-terminal domains (NTDs) of nsp2 preceding transmembrane helix 1 (TM1), which contains a protease domain (PLP2) and a long disordered loop (hypervariable region, HVR)^41^ (**Fig. 2a**, **Fig. 2h-j and Extended Data Fig. 4d-f**). The assembled pore complex comprises 12 copies each of nsp2 and nsp3, with an overall C3 symmetry, sharing a similar architecture with the CoV DMV pore complex. The EAV complex has an overall height of 150 Å and a width of 170 Å, spanning the double-membrane bilayer (**Fig. 2h**). 12 nsp2 on the outer membrane bilayer adopts C6 symmetry, forming two concentric alternating hexamer layers (**Fig. 2f, I**, dark and light blue), whereas 12 nsp3 on the inner membrane bilayer follows C3 symmetry (**Fig. 2g, j**). The overall membrane topologies of EAV nsp2 and nsp3 are highly consistent with those of their CoV counterparts, nsp3 and nsp4, respectively. Specifically, both the N– and C-terminal domains (NTD and CTD) of EAV nsp2 are exposed to the cytosol, whereas the EAV nsp3-CTD is oriented toward the DMV lumen, positioning their respective ectodomains reside within the intermembrane space (**Fig. 2a and Fig. 2e-m**). Notably, these EAV ectodomain pairs are substantially smaller than those of CoVs, resulting in a more compact overall structure on the membrane along the lateral axis (11.5 Å in EAV versus 83 Å in SARS-CoV-2; **Extended Data Fig. 5a-d**).

We then analyzed and compared the transmembrane domains (TMDs) of these two subunits. The TMD of EAV nsp2 is structurally different to its CoV nsp3. Specifically, EAV nsp2 contains four full transmembrane helices (TM1-4) and an additional half-spanning transmembrane helix (hTM). The first three helices (TM1-3) assemble into a unique three-helix bundle positioned on the peripheral side of the pore within the outer membrane, while the subsequent ectodomain, along with hTM and TM4, are situated proximal to the central pore (**Fig. 2l-m**). The second transmembrane protein, EAV nsp3, is a six-pass transmembrane protein and its TMD adopts distinct conformations in membrane to accommodate its local environment (**Fig. 2l-m**, yellow and orange**; Extended Data Fig. 5e-g**). EAV nsp3-TM1 is positioned separately from the main nsp3 TMD core (TM2-6), and ectodomain between TM1 and TM2-6 is arranged proximal to the pore axis. Within EAV nsp3 TMD core, its four-helix bundle (TM3-6) is relatively rigid whereas the TM2 displays a variable orientation (**Extended Data Fig. 5g**), recapitulating the dynamics of its CoV nsp4 (**Extended Data Fig. 5h-j**). The ectodomains of nsp2 and nsp3 are in direct contact within the intermembrane space. This remarkable structural conservation at the protomer level – despite vast sequence diversity – underpins the structural basis forming the universal architecture of DMV pore complex.

### Longitudinal and lateral interactions in EAV DMV pore complex

The DMV pore complex is stabilized by critical longitudinal and lateral interactions. The longitudinal heteromeric interactions locking the top nsp2 and bottom nsp3 layers occur primarily at the transmembrane helices near the central junction (**Fig. 3a-c**) and ectodomains (**Fig. 3b, d**). At the central junction, the twelve nsp3-TM1 helices interact intimately with the nsp2-hTM-TM4 bundles via a hydrophobic network, such as the aromatic stacking between nsp2-F380 and nsp3-Y8 (**Fig. 3c**). Within the DMV intermembrane space, twelve nsp2-nsp3 ectodomain pairs assemble into six radially arranged dimers of heterodimers. Interactions within these ectodomain interfaces are also predominantly driven by hydrophobic residues (**Fig. 3d**). Conversely, the lateral interactions are mainly driven by homomeric associations within the nsp2-Y1 domains (**Fig. 4**) and nsp3-CTDs (**Fig. 5**) located on opposite sides of the pore complex.

**Fig. 3.**
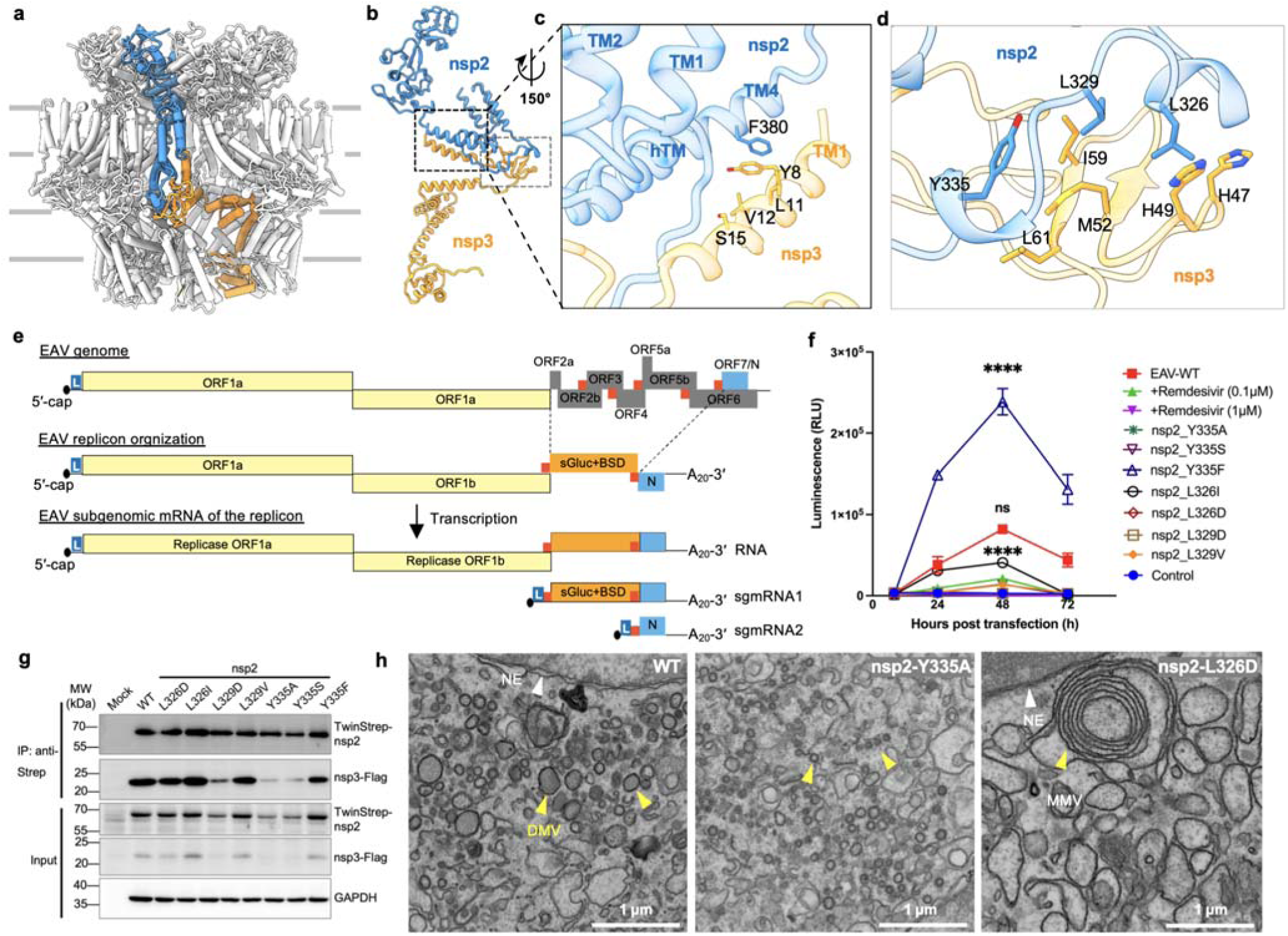
| Interaction between nsp2 and nsp3. **a**, Side view of the EAV DMV pore complex model with one pair of nsp2 (blue)-nsp3 (orange) protomers highlighted. **b**, Representative nsp2-nsp3 protomer, rotated around the z axis by 90° relative to **a**. **c**, Enlarged view of black boxed region in **b**, showing the interactions between nsp2 transmembrane domains and n p3-TM1, rotated approximately 150° relative to **b**. **d**, Enlarged view of the grey boxed region in b, indicating nsp2 and nsp3 ectodomains interface. **e,** Schematic of the EAV replicon constructed from the Bucyrus strain genome. The structural protein-coding region, except for ORF7 (N), was replaced with an sGluc-BSD reporter cassette. The replicon replicates and produces three subgenomic RNAs expressing sGluc and N**. f**, Luciferase activity in BHK-21 cells transfected with wild-type EAV replicon or mutants of key residues affecting hydrophobic interactions (blue residues in d) at indicated time points post-transfection. Statistical significance was determined using one-way analysis of variance (ANOVA) followed by Tukey’s multiple comparisons test using GraphPad Prism 9.0 (****P < 0.0001; ns, not significant). **g**, Co-immunoprecipitation assay showing disruption of hydrophobic interfaces between nsp2 and nsp3 ectodomains reduces nsp2-nsp3 binding. WT, wild type; Mock, cells transfected with empty vector. **h**, SEM micrographs of COS-7 cells transfected with wild-type or mutant nsp2-nsp3 constructs, showing defective DMV formation upon disruption of the ectodomain interface. NE, nuclear envelope (white arrowheads); DMVs, double-membrane vesicles (yellow arrowheads); MMV: multi-membrane vesicles. Scale bars are as indicated in the images. Scale bar, 1 µm. WT, wild type; Mock, cells transfected with empty vector.

**Fig. 4.**
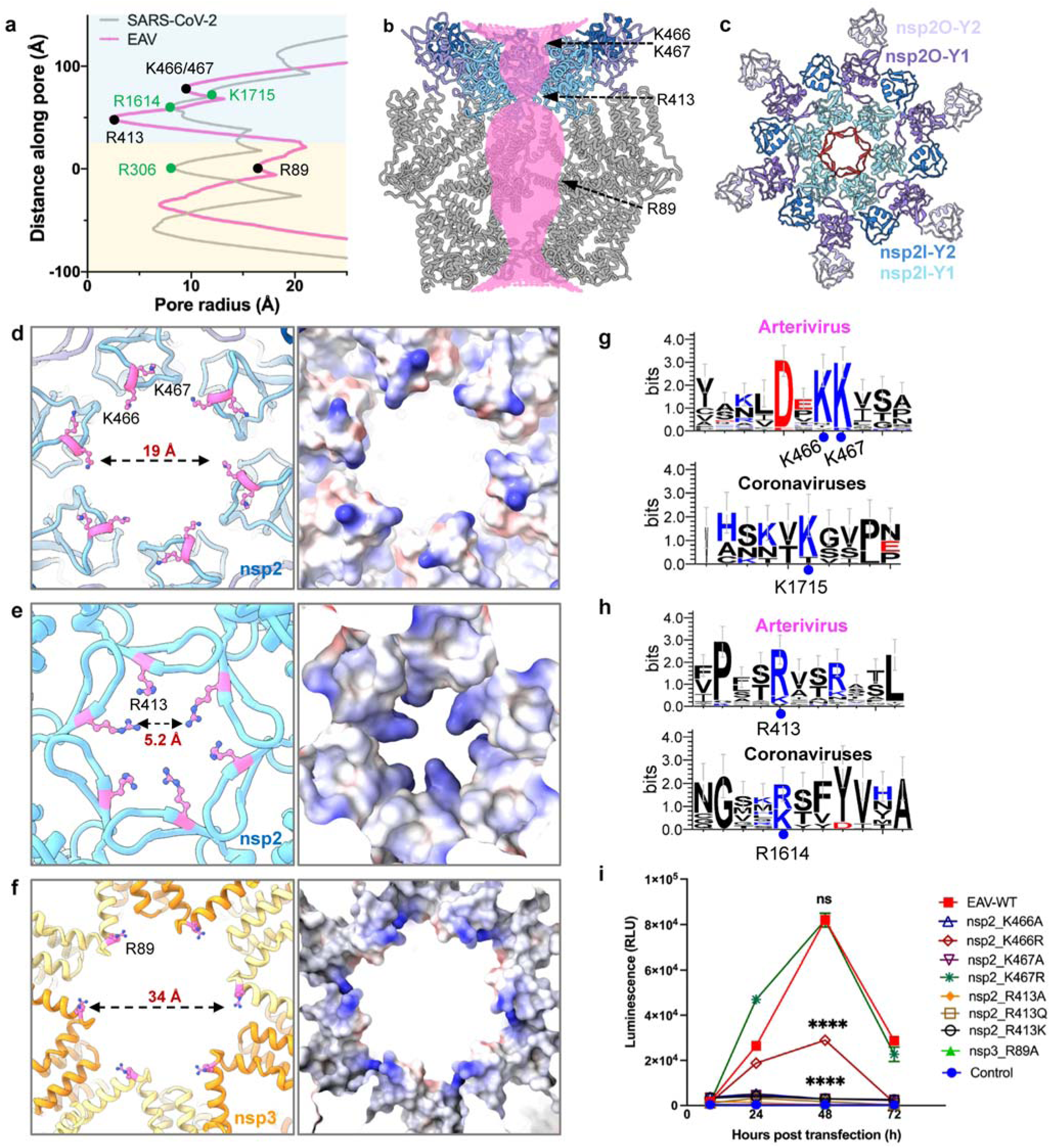
| Central channel of EAV DMV pore complex. **a**, Pore-radius profile of EAV DMV pore complex (pink) compared with that of SARS-CoV-2 (grey). Coloured dots indicate the residues at the three major constriction sites along the channel (black, EAV; green, SARS-CoV-2). Light blue and pale yellow shading indicate EAV nsp2 and nsp3 regions respectively, corresponding to coronavirus (CoV) nsp3 and nsp4. **b**, The central channel of the EAV DMV pore complex with the channel surface rendered in pink. Three major constriction sites are indicated by dashed lines. **c,** Top view of the EAV DMV pore complex showing the hexameric arrangement of inner nsp2 Y1/2 domains (nsp2I, cyan and blue) and outer nsp2 Y1/2 domains (nsp2O, purple and light purple). The central β-hairpin is coloured in red. **d–f,** Close-up top views of positively charged residues at the three major constriction sites lining the channel from top to bottom, with their corresponding electrostatic surface representations on the right: nsp2-K466/K467 (**d**), nsp2-R413 ring (**e**) and nsp3-R89 (**f**) at the membrane junction (corresponding to CoV nsp4-R306 ring). Distances represent pore radius in **a**. **g-h,** Sequence-logo conservation analysis of K466/K467 and R413 constriction sites in arteriviruses and equivalent positions in CoV, generated by WebLogo 3. Blue dots indicate the conserved residues. **i,** Luciferase activity in BHK-21 cells transfected with wild-type EAV replicon or positively charged pore mutants at indicated time points post-transfection. Statistical significance was determined using one-way analysis of variance (ANOVA) followed by Tukey’s multiple comparisons test using GraphPad Prism 9.0 (****P < 0.0001; ns, not significant).

**Figure 5.**
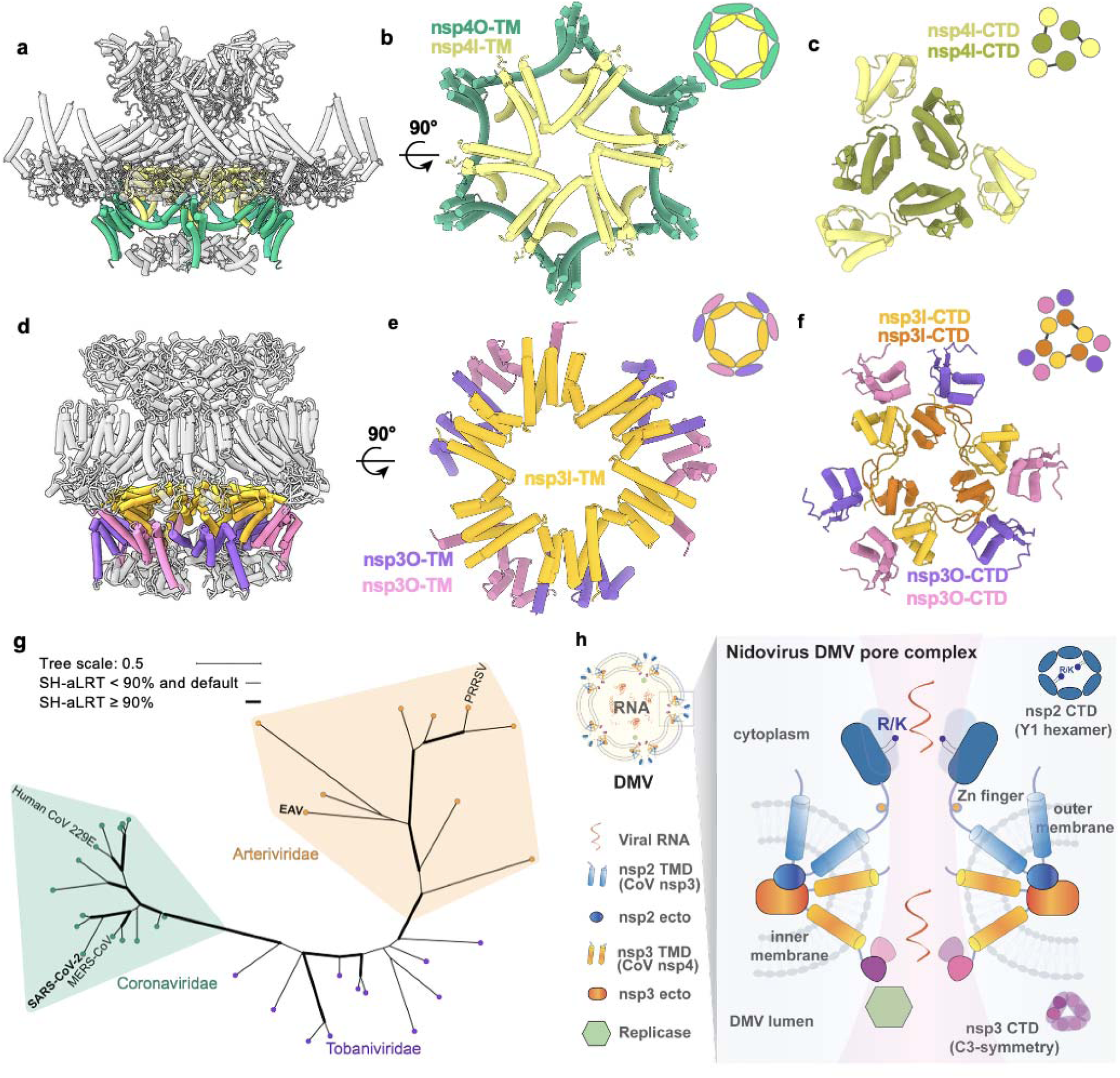
| Unified architectural features of DMV pore complexes from coronaviruses and arteriviruses. **a**, Side view of SARS-CoV-2 DMV pore complex (PDB:8YB7), with the inner nsp4 protomers (nsp4I) coloured in light yellow and outer nsp4 protomers (nsp4O) in green. **b,** Cross-section of the nsp4 transmembrane helices, with a schematic inset (top right) showing C6 symmetry for both inner and outer CoV nsp4 oligomers. **c,** Bottom view of the SARS-CoV-2 nsp4-CTD dimers assembled with C3 symmetry within the DMV pore complex. The nsp4–TD protomers are shown in yellow and olive; a schematic inset indicates the trimer of dimer configuration. **d**, Side view of the EAV DMV pore complex model. The inner nsp3 protomers (nsp3I) are shown in yellow and outer protomers (nsp3O) in pink and purple. **e**, Cross-section of the EAV nsp3 transmembrane helices, with a schematic inset (top right). The inner nsp3 oligomer exhibits C6 symmetry, whereas the outer nsp3 protomers show a C3 symmetry mismatch. **f**, Bottom view of the EAV DMV pore complex, with a schematic inset showing the C3-symmetric platform formed by four nsp3-CTDs (yellow, orange, pink and purple). The inner nsp3 (nsp3I) protomer is coloured orange to indicate the transition of the C3 symmetry mismatch from the transmembrane region towards the DMV lumen. **g**, A structural phylogenetic tree of nidovirus based on the conserved arterivirus nsp3 CTD/coronavirus nsp4 CTD fold. **h**, Model for nidovirus DMV pore formation.

To probe the functional significance of these interfaces in viral replication, we engineered an EAV virus genome derived replicon system. In this construct, ORFs 2–6 were replaced with a secreted luciferase reporter and a blasticidin S deaminase (BSD) resistance marker. All other viral genomic regions, including the replication machinery, the 5′ and 3′ untranslated regions (UTRs), the N gene, and cis-acting elements required for RNA replication, were retained unchanged (**Fig. 3e and Extended Data Fig. 6a**). We measured the luciferase reporter activity from the wild-type EAV replicon, which showed a progressive increase post transfection, reaching its highest level at 48 h, whereas the control group remained near background levels, indicating active RNA replication/transcription in this system. In contrast, remdesivir, a CoV RdRp inhibitor, caused a pronounced dose-dependent loss of signal, with 1 μM reducing luciferase activity almost completely (**Fig. 3f**). We subsequently evaluated these interface variants utilizing this engineered replicon. Targeted perturbation of the hydrophobic interfaces (Y335A, Y335S, L326D, L329D) severely diminished luciferase activity, whereas L329V mutation reduced and Y335F mutation enhanced the reporter signal, respectively (**Fig. 3f**). Co-immunoprecipitation assays further confirmed these interactions (**Fig. 3g**): disrupting interface hydrophobicity (L329D, Y335A, Y335S) abolished nsp2-nsp3 binding, correlating with diminished replicon activity. However, while the L326D mutant maintained the nsp2-nsp3 interaction, replicon function was nonetheless impaired.

To further evaluate whether these interface mutations specifically impaired DMV formation *in situ*, we conducted room-temperature scanning electron microscopy (RT-SEM) of COS-7 cells transfected with nsp2-3 constructs carrying variable mutations. Using the nuclear envelope (NE) as an internal reference (appearing as two parallel single membranes), vesicles with membranes visibly thicker than a single membrane are identified as DMVs (**Fig. 3h**). Consistent to previous study, the expression of WT nsp2–nsp3 remodel host membrane to form DMVs extensively (**Extended Data Fig. 7a-b**). All tested mutations negatively affected DMV formation efficiency, except Y335F. Charged substitutions (L326D and L329D) affected DMV formation as manifested by the substantially enlarged DMV and multiple-membranes vesicles (**Fig. 3h and Extended Data Fig. 7c-d**). In contrast, the Y335A mutation resulted in reduced DMV size, while Y335F exhibited a morphology very similar to that of the WT (**Extended Data Fig. 7e-f**). Together, the structural and functional analyses suggest the proper nsp2–nsp3 interactions helps to shape local DMV morphology and is critical to stabilize a replication-competent DMV pore architecture.

### Pore complex function and RNA transport channel

We next analyzed the central RNA transport channel. The EAV pore complex displays a distinct vase-like profile that differs substantially from its CoV counterpart (**Fig. 4a,b**; EAV, magenta; CoV, grey). There are three distinct constriction sites from the cytoplasm to the DMV lumen. At the cytoplasmic face, the Y1 domain (EAV nsp2) forms the narrowest constriction, measuring only ∼5 Å in diameter (**Fig. 4a,b**). In contrast, the middle constriction at the membrane junction, which serves as the narrowest point for RNA transport in CoV (17 Å in SARS-CoV-2 near nsp4-R306), is significantly wider in EAV at 34 Å (near nsp3-R89). Finally, the bottom luminal constriction formed by the nsp3-CTD domains is ∼15 Å in diameter, close to that of CoV (**Fig. 4b**, bottom).

On the cytoplasmic side, the EAV nsp2-Y1 domain structurally parallels the CoV Y1 domain by forming two concentric, alternating hexameric rings^42^ (**Fig. 4c** and **Extended Data Fig. 8a**). This domain contains a zinc-finger motif surrounded by positively charged residues that provide electrostatic complementarity to the lipid bilayer surface, analogous to CoV nsp3-Y1 (**Extended Data Fig. 8b-d**). The central channel of the homomeric Y1 hexamer is decorated with basic residues. Specifically, nsp2-K466 and K467 form a concentric lysine ring at the very top of the complex, while nsp2-R413 on a β-hairpin structure tightly lines the 5-Å constriction site, creating a highly positively charged bottleneck (**Fig. 4b-e and Extended Data Fig. 8c,d**).

To explore the functional relevance of these basic residues, we analyzed their sequence conservation across representative arteriviruses (**Fig. 4g-h** and **Extended Data Fig. 9**). AlphaFold3 predictions confirm that the structural arrangement of the Y1 hexamer and β-hairpin is highly conserved across the family (**Extended Data Fig. 8e**). This is remarkably analogous to the HIV capsid hexamer β-hairpin and Arginine-18 rings, which are essential for viral capsid assembly and dNTP import^43^. These observations raise the possibility that the translocation of single-stranded viral RNA likely requires outward movement of these arginines, potentially coupled with an iris-like opening of the β-hairpin structure. In addition to the basic residues in Y1 constrictions, we also found positively charged nsp3-R89 in the membrane junction (**Fig. 4f**), analogous to the CoV DMV pore (SARS-CoV-2 nsp4-R306). While EAV nsp3 R89 residue is not conserved across arteriviruses, the persistent presence of basic amino acids within this loop (between nsp3-TM2 and nsp3-TM3) points to a conserved functional role (**Extended Data Fig. 10**). The spatial localization of these basic residues in the arterivirus DMV pore corresponds precisely to CoV Y1 domains (**Fig. 4g-h**), indicating a functional convergence between arteriviruses and CoV.

We then introduced targeted mutations of these positively-charged residues along the pore to evaluate their contribution to replication via the replicon luciferase assay (**Fig. 4i**). Neutralization substitution (K466A, K467A) completely abolishes the replication whereas the K/R substitutions (K466R, K467R) mildly affected replication. The central R413 in the nsp2 Y1 β-hairpin is critical for replication, as R413A and R413Q completely abolish luciferase activity (**Fig. 4i**). Surprisingly, R413K also failed to support luciferase activity. The charge-neutralizing mutation (nsp3-R89A) noticeably diminished luciferase activity. These mutations do not affect nsp expression and DMV formation (**Extended Data Fig. 11**). Collectively, a more constricted, positively charged central pore defines the overall feature of the arterivirus DMV pore complex. The tight positively charged bottleneck indicates a channel tailored specifically for metabolite and single-stranded RNA transport, precluding the passage of double-stranded RNA or higher-order tertiary structures.

### Evolutionary conservation of nidovirus DMV pore complex

The structure of the EAV DMV pore complex provides crucial insights into structural conservation and divergence across the order *Nidovirales*. A major distinguishing feature between arterivirus and CoV DMV pore complexes is the spatial distribution of their symmetry mismatch. The TMDs of the CoV pore assemble with high C6 symmetry that extends throughout most of the complex (both the inner nsp4 and outer follows the C6 symmetry) (**Fig. 5a-b**), deviating only at the extreme luminal base by the inner nsp4-CTDs (nsp4I-CTD, **Fig. 5c**). In contrast, the EAV complex exhibits a pronounced C6-to-C3 symmetry mismatch. Specifically, the EAV nsp3 adopts an overall C3 symmetry, organized into a pseudo-C6-symmetric nsp3 inner hexamer (nsp3I) flanked by three outer nsp3 dimers (nsp3O, **Fig. 5d**). At the molecular level, this C3 symmetry is driven by the EAV nsp3-CTD (corresponding to CoV nsp4-CTD). The six inner nsp3I-CTDs pair into three antiparallel dimers stabilized by intermolecular β-strand exchange; each inner dimer is further flanked by two peripheral nsp3O-CTDs, together forming three equivalent I2O2 tetrameric modules around the pore (**Fig. 5f**). Together, these three tetramers coalesce to form a structural platform at the luminal base of the pore (**Fig. 5f**), with a local diameter of roughly 15 Å (**Fig. 4a-b**).

This luminal CTD assembly is critical for viral replication, as it may function as a highly specialized docking platform for the recruitment and attachment of the viral replicase complex within the DMV lumen^44^. To explore the structural conservation of this platform, we conducted homology search using EAV nsp3-CTD and SARS-CoV-2 nsp4-CTD structures against the predicted viral structural proteome. The resulting structure-based tree separated arteriviral and coronaviral proteins into distinct clusters (**Fig. 5g**). Tobaniviridae proteins can be identified by both arteriviral and coronaviral queries, occupying an intermediate position in the structure-based tree. Taken together, the structure-based tree reveals that nidovirus DMV luminal CTD domains share a conserved structural core but have undergone lineage-specific diversification in different nidoviral families. It indicates this luminal scaffold as a fundamentally conserved architectural principle across divergent nidoviruses.

## DISCUSSION

While the order *Nidovirales* is classically defined by its massive nsps and nested set of 3′-coterminal subgenomic mRNAs^12^, our work elevates this defining criteria to the ultrastructural level focusing on the replication organelle. Here, we propose the nidovirus replication organelle as a conserved, DMV-based compartment characterized by a unique, signature pore complex (**Fig. 5h**). By resolving the arterivirus DMV pore in atomic resolution, we provide compelling evidence that the transmembrane proteins preceding the replicase machinery not only commonly drive DMV formation across diverse nidoviral lineages, but also share a conserved molecular mechanism that dictates pore architecture (**Fig. 5**). The formation of this unique complex relies on the membrane-remodelling capacity of two nsps, driven by precise longitudinal and lateral interactions. We reveal that this overarching architecture is defined by several conserved structural hallmarks (**Fig. 5h**). The DMV pore complex is defined by a 12:12 stoichiometry of two transmembrane protein protomers organized into four stacked concentric rings on double-membrane, with a pronounced symmetry mismatch at the pore base. Functionally, the central pore is constricted by several sites including a cytoplasmic domain (the Y1 domain), junction with highly curved double-membrane in the middle and a trimeric DMV luminal domain in the bottom, featured with a central channel lined with basic residues critical for RNA transport (**Fig. 5h**).

Functionally, the viral genome can be simplified into two primary roles: nsps establish the replication organelle and dictate replicase function, while structural proteins package the RNA genome to form virion particles and mediate transmission^12^. DMV pore complexes emerge as a highly conserved feature across the order *Nidovirales*, contrasting with the vast diversity these viruses exhibit in genome size, virion morphology, and host tropism. Across nidoviruses, replication organelles exhibit significantly higher evolutionary conservation than the virion particles themselves, suggesting that these organelles may evolve partly independently of virion architecture^11^. The DMV pore complexes of arteriviruses and coronaviruses likely represent two structural extremes within the order: the EAV complex being a highly simplified variant, and the CoV complex representing a more elaborate, heavily decorated architecture. We attempted to computationally model the pore complexes of other Nidovirus families (*Mesoniviridae* and *Roniviridae*) but failed to obtain confident models. The scarcity of similar structures and the high diversity of these homologous sequences limit AlphaFold in predicting individual nsps confidently, and current algorithms struggle to accurately predict symmetry mismatches and corresponding conformational variations within multimeric assemblies. Future work will be required to fully map the evolutionary diversity and convergence of viral pore formation within Nidoviruses.

Finally, representing the smallest and simplest nidovirus DMV pore complex identified to date, the EAV complex serves as an elegant model for dissecting the core mechanisms of replication organelle biogenesis and viral RNA transport. Technically, the relatively small EAV genome enables rapid genetic manipulation and straightforward biochemical characterization while fully preserving the conserved features of the nidovirus pore. Our integrated cryo-ET and cryo-EM pipeline established here (**Extended Data Figs. 2 and 3**) provides a powerful blueprint for characterizing the replication organelle pore complexes of other positive-sense RNA viruses in their native membrane context^4^. By minimizing the structural complexity seen in larger viral families like Coronaviridae^8^, the EAV DMV system provides a unique opportunity to identify the minimal essential machinery required for DMV pore function and RNA transport. The insights from EAV pore complex highlights how structurally minimal viral systems can illuminate conserved targets across highly diverse virus orders, offering a foundation for future antiviral strategies development against Nidovirales, including coronaviruses.

## MATERIALS AND METHODS

### Plasmid Construction

The sequences encoding nsp2 (261-831aa) and nsp3 (832-1064aa) of Equine Arteritis Virus (UniProt code: P19811) were codon-optimized and cloned into a pcDNA3.1 vector with an N-terminal TwinStrep tag and a C-terminal Flag tag (pcDNA3.1-TwinStrep-nsp2-nsp3-Flag; synthesized by Tsingke). Site-directed mutagenesis of the full-length construct was performed by overlapping PCR.

### Expression and purification of EAV nsp2-nsp3 DMV

Human embryonic kidney 293F (HEK293F) cells (Expi293 Expression System Kit, A14635; Thermo Fisher Scientific) were cultured in OPM-293 CD05 medium (OPM Biosciences, 81075-001) supplemented with 100 U/ml penicillin and 100 U/ml streptomycin. To produce EAV nsp2-nsp3 DMVs, HEK293F suspension cultures at a density ∼3 × 10^6^ cells/ml were transfected with the EAV DMV plasmid using PEI MAX (Polysciences) at a DNA: PEI ratio of 1:4. At 20 h post-transfection, sodium butyrate was added to a final concentration of 2.5 mM. The cells were harvested at approximately 66 h (when cell viability reached 80-85%) post-transfection by centrifugation at 2,000 × g and washed with 1× phosphate-buffered saline (PBS). The resulting cell pellets were flash frozen in liquid nitrogen and stored at −80 °C until further use.

To preserve the integrity of EAV DMVs, all purification steps were performed on ice or at 4 °C as described previously^15^. Frozen cell pellets from 2L cell culture were resuspended in 200mL hypotonic buffer (20 mM HEPES, pH 7.5, 1.5 mM MgCl_₂_, 10 mM KCl) with protease inhibitor cocktail (Sigma-Aldrich, P8340) and GENIUS nuclease (ACROBiosystems, BEE-N3116). After incubation at 4 °C for 1 h, the cell suspension were homogenized on ice with a 40 mL Dounce homogenizer (Thomas Scientific) by 40 strokes, followed by sonication (2 s on, 4 s off, 30% power) for 60 cycles by a probe sonicator (Branson Digital Sonifier SFX 550). The homogenate was centrifuged at 4,000 × g for 20 min to remove the remaining cell debris. The supernatant was supplemented with 150 mM NaCl and incubated with 1ml Strep-Tactin® XT 4Flow resin (2ml 50% suspension) (IBA LifeSciences, 2-5010-025) pre-equilibrated with equilibration buffer (50 mM HEPES, pH 7.5, 150 mM NaCl, 1 mM EDTA, 5% glycerol) for 3 h at 4 °C in a gravity column. The flow-through was discarded and the resin was washed with 50 column volume (CV) washing buffer (50 mM HEPES, pH 7.5, 500 mM NaCl, 1 mM EDTA, 5% glycerol) and 10 CV equilibration buffer. Bound vesicles were eluted by elution buffer (50 mM HEPES, pH 7.5, 150 mM NaCl, 1 mM EDTA, 5% glycerol and 50 mM biotin) for 4 times, with a 10 min incubation period for each. All elution fractions were pooled together, transferred to an ultracentrifuge tube, and centrifugated at 200,000 × g for 1 h using a Thermo Scientific Sorvall WX ultracentrifuge equipped with a TH-641 swing-bucket rotor. The resulting pellets were resuspended in 10 μl STE buffer (10 mM Tris-HCl pH 8.0, 150 mM NaCl and 1 mM EDTA), and used for cryo-EM grids preparation immediately.

### Co-immunoprecipitation and western blot analysis

HEK293T cells grown on 10-cm dishes at ∼75% confluence were transfected with wild-type or mutant pcDNA3.1-TwinStrep-nsp2-nsp3-Flag plasmids. Transfection were performed using PEI MAX with 10 μg plasmid DNA per dish at a DNA:PEI ratio of 1:3. At 42 h post-transfection, cells were washed with PBS and harvested by centrifugation at 2,000 × g for 10 min, followed by twice more washes with PBS. Cell pellets from each 10-cm dish were lysed in 800 μl of ice-cold lysis buffer (50 mM Tris-HCl, pH 7.5, 150 mM NaCl, 5% glycerol and 1% Triton X-100) supplemented with protease inhibitor cocktail (Sigma-Aldrich, P8340) and GENIUS Nuclease (ACROBiosystems, BEE-N3116). Lysates were incubated on ice for 30 min and sonicated on ice using a Branson Digital Sonifier SFX 550 probe sonicator at 30% amplitude (2s on/5s off) for a total sonication time of 16 s per sample. Cell debris was removed by centrifugation at 12,000 × g for 20 min at 4 °C.

As an input control, an aliquot of the clarified lysate (40 μl) was mixed with 10 μl 4× SDS sample buffer (0.2 M Tris-HCl, pH 6.5, 0.4 M dithiothreitol, 8% SDS, 6 mM bromophenol blue and 4.3 mM glycerol), and heated at 75 °C for 10 min. The remaining lysate was incubated with 60 μ Strep-TactinXT 4Flow resin (pre-equilibrated as 50% resin suspension in lysis buffer; IBA LifeSciences, 2-5010-025) for 2 h at 4 °C with gentle rotation. The resin was washed three times with 1 ml ice-cold lysis buffer, and bound proteins were eluted with 90 μl 2× SDS sample buffer and heated at 75 °C for 10 min. The 10 μl of input and 15 μl of elution samples were loaded for western blot analysis. Proteins were separated on 12% SDS–polyacrylamide gels at 160 V for 90 min and transferred to PVDF membranes (Millipore, IPVH00010) at 220 mA for 2.5 h. Membranes were blocked in 5% non-fat milk (Santa Cruz Biotechnology, sc-2325) prepared in TBST buffer (50 mM Tris-HCl, pH 7.4, 150 mM NaCl and 0.1% Tween-20) for 2 h at room temperature. Following three 5-min washes with TBST, membranes were incubated overnight with primary antibodies diluted in TBST containing 0.05% non-fat milk. The primary antibodies included: mouse anti-Strep antibody (IBA, 2-1507-001; 1:5,000), mouse anti-Flag/DYKDDDDK monoclonal antibody (Thermo Scientific, MA1-91878; 1:1,000) and mouse anti-GAPDH antibody (Santa Cruz Biotechnology, sc-47724; 1:5,000). Membranes were then washed three times with TBST and incubated for 1 h at room temperature with HRP-conjugated anti-mouse IgG secondary antibody (Cell Signaling Technology, 7076S; 1:3,000) in TBST containing 0.05% non-fat milk. After three final washes with TBST, signals were developed using chemiluminescent substrate (Thermo Scientific, 34095) and visualized with a ChemiDoc MP Imaging System (Bio-Rad).

### Cryo-EM/ET grid preparation of isolated EAV nsp2-nsp3 DMVs

To disperse large aggregates and achieve thinner ice on the cryo-EM grids, purified DMV samples were sonicated in water bath until the suspension became visually transparent. For cryo-ET, 6-nm bovine serum albumin-coated gold fiducial beads (Aurion, 206.033) were added to the sample prior to plunge freezing; these beads were omitted for samples intended for single particle cryo-EM. A 3.5 µl of DMV samples was applied to freshly glow-discharged lacey carbon grids (Agar Scientific, AGS166-3). The grids were blotted for 3.5s with a blot force of 0 at 4 °C with 100% humidity before being plunge-frozen in liquid ethane using a Vitrobot Mark IV (Thermo Fisher Scientific).

### Cryo-ET data collection and STA of isolated EAV nsp2-nsp3 DMVs

Tilt series of EAV DMVs were acquired using a Titan Krios G4 (Thermo Fisher Scientific) operating at 300 kV, equipped with a Falcon 4i camera and a Selectris energy filter (20 eV slit). Data acquisition was performed using the PACE-tomo script^45^ within SerialEM^46^, allowing a maximum image–beam shift of 15 µm when adding acquisition points. A dose-symmetric scheme (group of 2) was employed over a tilt range of ± 48° in 3° increments. Images were recorded at a nominal magnification of 81,000 × (pixel size: 1.571 Å) with a total exposure dose of 3 e^−^/Å² per tilt and a defocus range of −1.5 to −4.5 µm. A total of 1,708 tilt series were collected over several sessions on multiple grids from two independent DMV purifications. Acquisition locations were manually selected from medium-magnification montages at 4,800 ×, where DMVs appeared as slightly electron-dense vesicles. Among these, vesicles containing visible DMV pore complexes were prioritized for data collection. The detailed data collection parameters are listed in **Extended Data Table 1**.

Raw movie frames in EER format were motion-corrected and gain normalized using relion_motioncor implementation in RELION 4.0^47^. Blank images were removed based on average image intensity using the clip command in imod/4.11.24. Tilt-series alignment was performed using batchruntomo with the Python script (tomo_toolbox.py: https://github.com/ffyr2w/cet_toolbox). CTF parameters were estimated in emClarity (v1.6.1.0)^48^. To generate an initial template, 300 particles were manually picked from 19 reconstructed tomograms using the ArtiaX^49^ plugin in ChimeraX^50^, with their z-orientations predefined as perpendicular to membrane. These particles (bin4) were subjected to 3D classification and refinement in RELION 4.0. Although the precise symmetry of the map could not be determined at this stage, both C1 and C6 symmetries were evaluated during data processing. The C6-symmetrized density map was further refined with 2×– binned particles and used as the template to pick all the particles via template-matching in emClarity (v1.6.1.0). A total of 378,650 particles were picked from 1,295 manually inspected tomograms containing DMVs and these particles were exported to RELION 4.0, for further 3D classification and 3D refinement. Particles from five good classes were pooled together and refined with 4×-, 2×-and 1×-binned pseudosubtomograms with C6 symmetry applied. Following further tomo frame alignment, a final reconstruction of the pore complex was obtained at an overall resolution of 7.5□Å (**Extended Data Table 1**; **Extended Data Fig. 2**).

### Cryo-EM data collection and SPA of isolated EAV nsp2-nsp3 DMVs

The single particle cryo-EM dataset of EAV DMVs was acquired using the same microscope and the acquisition positions were manually selected in SerialEM with PACE-tomo script as described for the above cryoET dataset. Micrographs were recorded at a nominal magnification of 130,000 <u>×</u> (pixel size: 0.9557 Å) with a total dose of 40 e−/Å² and a defocus range of −0.5 to −1.5 μm. In total, 33,273 movies were collected over several sessions on multiple grids prepared from two independent DMV purifications (**Extended Data Table 1; Extended Data Fig. 3**).

Raw micrographs in EER format were motion-corrected and gain normalized using relion_motioncor implementation in RELION 4.0.The motion-corrected micrographs were binned by a factor of 2 (final pixel size: 1.9114 Å) using binvol command in IMOD/4.11.24, followed by CTF estimation using CTFFIND4^51^ within *cis*TEM (2.0.0-alapha-328)^52^ All micrographs were manually inspected to discard poor-quality images or those lacking DMVs, resulting in a curated dataset of 21,185 micrographs for the following 2D template matching in *cis*TEM (2.0.0-alapha-328)^53^. For the first round of particle picking, the 7.5 Å cryo-ET reconstruction was rescaled and used as a template on a subset of 10,823 micrographs. Template matching searches were performed with C6 symmetry, utilizing an out-of-plane angular step of 1.5°, an in-plane angular step of 1.0° and a high-resolution limit of 4.5 Å. Particle coordinates were refined in *cis*TEM using the refine_template script, and the detected target parameters (including particle positions, local defocus values and Euler angles) were extracted using prepare_stack_matchtemplate. A total of 989,684 particles were extracted by applying a cross-correlation peak-height threshold of 6.5, and their coordinates were exported as a STAR file for subsequent processing in RELION. Following one round of 2D classification, obvious false positives, such as carbon area, single membranes and duplicated particles (distance less than 50 Å) were removed, leaving 742,075 particles for 3D classification. These particles were subjected to two rounds of 3D classification applying C6 symmetry, and classes corresponding to single membranes, double membranes without junctions, or particles lacking membrane-curvature features were discarded. The remaining 194,067 particles were subjected to 3D refinement applying C6 symmetry at bin2 and subsequently bin1 (**Extended Data Fig. 3a-d**).

The resulting 4.9 Å map was rescaled to bin2 and used as an improved template for a second round of template matching on the complete dataset (21,185 micrographs) with the same parameters used in the first round. The peak threshold for particle extraction was set to 6.5, yielding 1,298,306 particles. Particles were firstly extracted in RELION at 2 × binning and subjected to 2D classification to discard obvious false positives. Subsequently, three rounds of 3D classification were performed with the 2×-binned particles with C6 symmetry and all bad particles were discarded. Classes displaying well-defined structural features, comprising 597,183 particles, were selected, re-extracted without binning, and subjected to 3D refinement with C6 symmetry. After one round of Bayesian polishing followed by 3D refinement with C6 symmetry, 3D classification was performed using an angular sampling interval of 0.9° with local angular search (sigma_ang 2). From the ten classes, one class containing 96,354 particles (16.13% of the input particles) showed clear secondary-structure features and was selected for further 3D refinement with C6 symmetry. To improve the density of the nsp3 C-terminal domain, focused 3D classification was performed using a mask covering the nsp3 region, including the nsp3 C-terminal domain and the transmembrane regions within the inner membranes. C3 symmetry was imposed based on the previously reported C3-symmetric organization of the SARS-CoV nsp4 C-terminal domain. Two major classes with clear structural features in the nsp3 C-terminal domain were selected, containing 37,052 (38.13%) and 39,060 (40.18%) particles, respectively. Particles from one class were rotated by 180° to align with the other class, and the combined particles were subjected to further 3D local refinement, yielding a reconstruction at approximately 4 Å resolution. Following this refinement, Bayesian polishing and CTF refinement, including beam-tilt correction, anisotropic magnification correction, per-particle CTF refinement and per-micrograph astigmatism refinement, were performed. A subsequent iteration of Bayesian polishing and 3D refinement with imposed C3 symmetry improved the resolution to 3.58 Å. A final masked 3D refinement with C3 symmetry achieved a reconstruction with an overall resolution of approximately 3.0 Å (**Extended Data Fig. 3e-j**).

### Model building and validation

An initial atomic model of EAV nsp2-nsp3 complex was generated de novo using CryoAtom^54^ from the 3.0 Å resolution C3-symmetry map. Missing regions of the initial model were manually built in Coot^55^. The complete model (nsp2 262-571aa, nsp3 1-233aa) was subsequently refined using ISOLDE^56^ implemented in ChimeraX. The resulting complex structure was subject to iterative cycles of real-space refinement in Phenix^57^ and Coot. The final model was validated using Phenix with MolProbity^58^, and the fitness of model to density was measured by Q-score^59^. The refinement parameters and validation statistics are summarized in **Extended Data Table 1**. The models were presented in ChimeraX.

### Scanning electron microscopy

#### SEM Sample preparation and en bloc staining

Monolayers of COS-7 cells cultured on coverslips were transfected with wild-type or mutant pcDNA3.1-TwinStrep-nsp2-nsp3-Flag constructs using Lipofectamine 3000 (cat. no. L3000015, Thermo Fisher Scientific, USA). At 36 h post-transfection, cells were washed three times with pre-warmed (37 °C) serum-free DMEM. Primary fixation was performed using pre-warmed 2.5% glutaraldehyde (diluted from 50% EM grade stock, cat. no. 16320; Electron Microscopy Sciences (EMS), USA) in 0.1 M sodium cacodylate buffer (pH 7.2-7.4; prepared from sodium cacodylate trihydrate, cat. no. 12310; EMS) for 10 min at room temperature (RT). This was followed by a solution exchange with fresh 2.5% glutaraldehyde for further fixation overnight at 4 °C. Following primary fixation, the specimens were rinsed five times (5 min each) with fresh 0.1 M cacodylate buffer. To enhance lipid and membrane contrast, an OTO (osmium-thiocarbohydrazide-osmium) post-fixation protocol^60^ was employed. Briefly, samples were incubated in a solution of 2% osmium tetroxide (OsO_₄_; diluted from 4% aqueous solution, cat. no. 19190; Electron Microscopy Sciences) and 1.5% potassium ferricyanide in 0.1 M cacodylate buffer for 1 h on ice. After five 5-min washes with ice-cold double-distilled water (ddH_₂_O), specimens were incubated in 1% thiocarbohydrazide (TCH; cat. no. 21900; Electron Microscopy Sciences) in ddH_₂_O for 20 min at RT. The samples were washed again with ddH_₂_O prior to a second osmication step using 2% OsO_₄_ in ddH_₂_O for 30 min at RT. After extensive washing with ddH_₂_O, a final *en bloc* staining was performed using 2% neodymium (III) acetate (cat. no. 325805, Sigma-Aldrich, USA) overnight at 4 °C.

#### Dehydration and embedding

Following overnight *en bloc* staining, residual neodymium was removed via five 5-min washes with ice-cold ddH_₂_O.The specimens were then dehydrated on ice through a graded series of ice-cold ethanol (Fisher Scientific, UK): 30%, 50%, 70%, 85%, 95% and 100% (twice) for 10 min per step. A final 10-min dehydration step was performed using 100% ethanol at RT, after which the ethanol was replaced with 100% acetone. The dehydrated specimens were infiltrated with EMbed 812 resin (cat. no. 14121, Electron Microscopy Sciences, USA) via sequential incubations in 1:1 and 3:1 (v/v) resin: acetone mixtures for 2 h each. This was followed by two changes of pure resin for 2 h each. Finally, BEEM capsules (cat. no. 133-P; Ted Pella, Inc., USA) filled with fresh resin were inverted onto the coverslips, and the resin was polymerized at 60 °C for 48 h.

#### Sectioning and imaging

Following polymerization, coverslips were detached, and the resin blocks were trimmed with a razor blade to expose the region of interest. Sections (∼200 nm thick) were cut using an UC Enuity ultramicrotome (Leica Microsystems, Germany) equipped with a diamond histo-knife (Diatome Ltd., Switzerland) and subsequently collected onto conductive silicon wafers. Sections were examined using a field-emission scanning electron microscope (FE-SEM; ZEISS GeminiSEM 360, Germany). Micrographs were acquired under high vacuum conditions at an accelerating voltage of 5 kV using an Inlens SE detector. SEM images of cellular double-membrane vesicles (DMVs) and endoplasmic reticulum (ER) clusters were acquired at a magnification of 15,000 ×. Ten representative micrographs were acquired per sample (pooled from two independent biological replicates, n = 2). To ensure a comprehensive evaluation, an initial systematic screening was performed across four to five sections. This extensive observation encompassed more than a thousand cells, allowing us to accurately identify and categorize the dominant structural features. Subsequently, to minimize sampling bias, three to four distinct, non-overlapping fields of view were imaged to document each specific morphological feature.

### Construction of EAV replicon and full-length infectious clone

A plasmid encoding the full-length complementary DNA (cDNA) of the Bucyrus strain (VBS; ATCC VR-796) of Equine Arteritis Virus (EAV; Taxonomy ID: 299386) was synthesized *de novo* (Tsingke Biotechnology Co., Ltd., Chengdu, China). To generate a self-replicating RNA replicon suitable for high-throughput screening, the genomic regions encoding the major structural proteins (M, E, GP2b, GP3, GP4, and GP5) were precisely replaced with a bicistronic reporter-selectable marker cassette^61^, which gifted from Professor Zhigang Yi at Fudan University (Shanghai, China). While the nucleocapsid (N) gene was retained in its native genomic position to RNA replication. The inserted cassette consists of: (i) a codon-optimized secreted Gaussia luciferase (sGluc) gene under the control of a T7 promoter; (ii) a foot-and-mouth disease virus (FMDV) 2A autoproteolytic peptide (sequence: NFDLLKLAGDVESNPGP); and (iii) a blasticidin S deaminase (BSD) resistance gene, which serves as a selectable marker for mammalian cell culture. The 3′ untranslated region (3′ UTR) was stabilized by the addition of a 20-nucleotide polyadenylation signal (polyA20). A schematic representation of the recombinant EAV replicon construct is provided in **Fig. 3d and Extended Data Fig. 6**.

Site-directed mutagenesis was performed on the full-length replicon plasmid using overlapping polymerase chain reaction (PCR) to introduce specific amino acid substitutions in two distinct non-structural proteins: nsp2 (L326D, L329D, Y335F, Y335A, R413A, R413K, R413Q, K466A, K466R, K467A, K467R) and nsp3 (R89A). All mutagenic primers were synthesized by SINBIO (HongKong, China). Plasmid integrity and mutation accuracy were verified by Sanger sequencing (SINBIO, HongKong, China). Only clones with correct nucleotide sequences were selected for downstream applications.

The full-length infectious clone was generated by restoring the deleted structural protein genes into the replicon backbone. Specifically, a DNA fragment encoding the complete set of EAV structural proteins (M, E, GP2b, GP3, GP4, and GP5) was synthesized (Tsingke Biotechnology Co., Ltd.). This fragment was used to replace the *sGluc-2A-BSD* reporter-selectable marker cassette in the parental replicon plasmid. Plasmid integrity was confirmed by Sanger sequencing (SINBIO, HongKong, China).

### Replicon RNA in vitro transcription, purification and quantification

For *in vitro* transcription, 5 µg of purified replicon plasmid DNA were linearized by XbaI (New England Biolabs, USA; Cat. No. R0145S) at 37°C for 2 h. Linearization efficiency was confirmed by agarose gel electrophoresis (1% w/v, 1× TAE buffer). One microgram of the linearized DNA template was used for *in vitro* transcription using the mMESSAGE mMACHINE T7 Transcription Kit (Thermo Fisher Scientific, USA; Cat. No. AM1344). The 20 µl reaction mixture containing 1 µg linearized DNA template, 2 µl 10× Reaction Buffer, 10 µl 2× NTPs mixture and 2 µl T7 RNA Polymerase was incubated at 37°C for 2 h in a water bath. Following transcription, the reaction was treated with 2 U of TURBO DNase (supplied with transcription kit) at 37°C for 15 min to remove residual DNA template.

Transcribed RNA was purified using the RNeasy Mini Elute Cleanup Kit (Qiagen, Germany; Cat. No. 74106) according to the manufacturer’s protocol. Briefly, the transcription reaction was mixed with 350 µl RLT buffer containing 1% β-mercaptoethanol, passed through a spin column, and washed twice with RW1 buffer and once with RPE buffer. RNA was eluted in 20 µl of nuclease-free water (supplied with kit) by centrifugation at 14,000 × g for 2 min. RNA concentration and purity were determined by measuring the absorbance at 260 nm and 280 nm using a NanoDrop 2000 spectrophotometer (Thermo Fisher Scientific). RNA integrity was assessed by denaturing agarose gel electrophoresis using 1% (w/v) Agarose in 1 × MOPS buffer (40 mM MOPS, 10 mM NaOAc, 1 mM EDTA, pH 7.0) containing 6.3% formaldehyde. The gels were stained with SYBR™ Green II RNA Gel Stain (Invitrogen, Cat. No. S7564) and visualized using a ChemiDoc MP imaging system (Bio-Rad). The transcribed RNA was aliquoted and stored at –80°C in aliquots. To maintain RNA integrity, repeated freeze-thaw cycles were strictly avoided.

### Luciferase assays of EAV replicons

BHK-21 cells (Baby Hamster Kidney cells, ATCC CCL-10) were maintained in Dulbecco’s Modified Eagle Medium (DMEM) supplemented with 10% (v/v) fetal bovine serum (FBS; Gibco, Cat. No. 10270106), 100□U/ml penicillin, and 100□U/ml streptomycin (Gibco, Cat. No. 15140122) at 37°C in a humidified 5% CO_₂_ incubator. For transfection, BHK-21 cells were seeded at a density of 1.8 × 10□ cells per well in 24-well tissue culture plates and cultured overnight to achieve reach ∼70–80% confluency. Transfection of *in vitro* transcribed replicon RNA (0.5 µg per well) was performed using the TransIT-mRNA Transfection Kit (Mirus Bio LLC; Cat. No. MIR 2255). Briefly, 0.5 µg RNA was complexed with 1 µl TransIT-mRNA Reagent and 1 µl mRNA Boost Reagent in 50 µl Opti-MEM I Reduced Serum Medium (Gibco, Cat. No. 31985070) for 10 min at room temperature. The RNA-lipid complex was then added dropwise to the cells, followed by incubation at 37°C in 5% CO_₂_ incubator. At 8 h post-transfection, the transfection medium was replaced with fresh complete DMEM containing 10% FBS. For the Remdesivir treatment group, cells were pretreated with Remdesivir (GS-5734; MedChemExpress, Cat. No. HY-101769) at a final concentration of 100 nM for 4 hours prior to RNA transfection. Following transfection, the cell medium was replaced at 8 hours post-transfection with fresh DMEM medium supplemented with 10% (v/v) FBS.

At predetermined time points post-transfection (8, 24, 48, and 72 hpt), 20 µl of cell culture supernatant was collected from each well and mixed with an equal volume of 2× Passive Lysis Buffer (Promega, Cat. No. E2820). The clarified supernatant was transferred to a white 96-well plate, and luciferase activity was measured using the Renilla Luciferase Assay System (Promega, Cat. No. E2820). Upon the addition of 50 µl of Luciferase Assay Reagent (LAR) per well, luminescence was recorded as relative light units (RLU) using a 3-s integration period with a 0.5 s delay of 0.5 seconds. The emission wavelength was ∼480 nm. All experiments were performed in triplicate (n = 3). Data are presented as mean ± standard deviation (SD). Statistical significance was determined using one-way analysis of variance (ANOVA) followed by Tukey’s multiple comparisons test using GraphPad Prism 9.0 (GraphPad Software, La Jolla, CA, USA). A p-value < 0.05 was considered statistically significant.

### EAV virus rescue in Huh7 cells

Human hepatoma Huh7 cells (JCRB0403) were maintained in DMEM supplemented with 10% (v/v) FBS, 100 U/ml penicillin, and 100 U/ml streptomycin. at 37°C in a 5% CO2 incubator. Infectious virus was rescued by electroporation of *in vitro*-transcribed genomic RNA into Huh7 cells. Briefly, a 90% confluent T75 flask of Huh7 cells was trypsinized, washed twice with ice-cold Phosphate-Buffered Saline (PBS), and resuspended in cold PBS at a density of 1.5 × 10 cells/ml. A 400 µl aliquot of the cell suspension (5 × 10^6^ cells) was mixed with 5 µg of full-length EAV genomic RNA in a 0.4 cm electroporation cuvette (Bio-Rad, Cat. No. 1652081). Electroporation was performed using a Gene Pulser II (Bio-Rad) set to 270 V, 950 µF, and infinite resistance. After electroporation, cells were rested at room temperature for 10 min, then transferred to a T25 flask containing 5 ml of pre-warmed complete growth medium. The medium was replaced 24 h post-electroporation. At 5 days post-electroporation, the culture supernatant, containing the rescued virus, was harvested, clarified by centrifugation at 2,000 × g for 10 minutes, filtered through a 0.45 µm PES membrane filter (Millipore, Cat. No. SLHV033RS), aliquoted, and stored at –80°C. All work with live EAV was conducted in a Biosafety Level 2 (BSL-2) facility.

### EAV Virus Propagation and Titration

To generate high-titer virus stocks, filtered EAV supernatant was used to inoculate fresh Huh7 cells in five T75 flasks at a low multiplicity of infection (MOI ≈ 0.01). The supernatant was collected 3-4 days post-infection when extensive cytopathic effect (CPE) was observed. The pooled supernatant was clarified by centrifugation and filtration (0.45 µm). The concentrated virus stock was aliquoted and stored at –80°C. Virus infectious titer was determined by the 50% tissue culture infectious dose (TCID_50_) assay on Huh7 cells. Briefly, ten-fold serial dilutions of the virus stock were prepared in serum-free DMEM. 100 µl of each dilution was added to eight replicate wells of a 96-well plate containing Huh7 cells at 80% confluency. After 2 h of adsorption at 37°C, the inoculum was removed, and 100 µl of complete medium was added. The plate was incubated for 4-5 days, and wells were scored for the presence or absence of CPE. The TCID_₅₀_/ml was calculated using the Reed-Muench method.

### EAV Inactivation and Validation

EAV in infected cell culture supernatants was inactivated using formaldehyde fixation. Formaldehyde (Sigma-Aldrich, Cat. No. 252549) was added directly to the supernatant to achieve a final concentration of 3.7% (v/v). The mixture was incubated at room temperature for the indicated durations (5, 10, 15, 30, or 60 minutes). The reaction was terminated by removing the fixative and washing the treated sample three times with PBS. The efficacy of inactivation was confirmed by two independent methods: RT-qPCR for the detection of residual viral genomic RNA, and a plaque assay for the detection of infectious virus. RT-PCR: Fresh Huh7 cells in 24-well plates were inoculated with 100 µL of inactivated supernatant. Total RNA was extracted from the cell culture supernatant at 24 and 72 hours post-inoculation (hpi) using the RNeasy Mini Kit (Qiagen, Cat. No. 74104). Residual viral genomic RNA was detected by one-step RT-qPCR using the One Step TB Green PrimeScript RT-PCR Kit II (Takara, Cat. No. RR086A). EAV nucleocapsid (N) gene-specific primers were used: EAV-N-F: 5’-ATGTTTGGTCAGATGCGGGT-3’; EAV-N-R: 5’-CCAACTGACGGTGTACGTGA-3’. The reaction conditions were: 42°C for 5 min, 95°C for 10 s; followed by 40 cycles of 95°C for 5 s and 60°C for 30 s. A standard curve generated from a plasmid of known copy number was used for absolute quantification. Complete inactivation was defined as Cq values indistinguishable from the mock-infected control. For plaque assay, Huh7 cells in 6-well plates were inoculated with 200 µL of inactivated supernatant for 2 hours. The inoculum was then removed, and cells were overlaid with 1 mL of a 1:1 mixture of 2× DMEM and 1.2% low-melting-point agarose (Invitrogen, Cat. No. 16520100). After solidification at room temperature for 20 minutes, plates were incubated at 37°C for 48-72 hours. Cells were then fixed overnight with 4% paraformaldehyde, the overlay was removed, and the cell monolayer was stained with 0.5% crystal violet for 10 minutes. The absence of plaque formation indicated complete viral inactivation.

### In situ cryo-ET of EAV-infected cell

#### EAV infection, micropatterning and cryo-ET grid preparation

For cryo-electron tomography (cryo-ET) sample preparation, Quantifoil Holey Carbon Film R 2/2 on Gold 200 mesh Grids (Quantifoil, N1-C16nAu20-01) were glow discharged for 60 s at 15 mA with PELCO easiGlowTM Glow Discharge Cleaning System and transferred carbon-side up to a coverslip for passivation. The PDMS stencils (Alvéole) were used to cover grids while leaving a grid-sized opening for stabilization on the coverslip. A droplet of poly-L-lysine (PLL; Sigma-Aldrich, Cat. No. P8920, diluted to 1x) was applied and incubated for 30 minutes. After washing with 0.1M HEPES at pH 8.3, the surface was passivated with 100 mg/mL mPEG-SVA (Laysan Bio) for 1 hour, followed by washing five times in 1x PBS at pH 7.4. The PLPP gel solution (Alvéole) diluted in 70% ethanol was then applied on grids, followed by completely drying. The coverslip then was placed on the microscope with the Primo 2 micropatterning unit equipped a UV pulsed laser source of 355 nm. The patterns were created on EM grids by UV laser with 40 mJ/mm dose and 25 µm diameter. The micropatterned surface was functionalized by incubating with human plasma fibronectin (Corning, Cat. No. 356008, 20 µg/mL in PBS) for 30 minutes at room temperature. After washing with 1x PBS at pH 7.4, the grids were exposed to UV light for 1 hour for sterilization.

The micropatterned grids were placed on a 6-well plate for cell seeding. Human hepatoma Huh7 cells (JCRB0403) were maintained in DMEM supplemented with 10% (v/v) FBS, 100 U/ml penicillin, and 100 U/ml streptomycin. at 37°C in a 5% CO2 incubator. After trypsinization, a volume of 10 µL of a 3 × 10^5^ cells/mL huh-7 cells were seeded on the micropatterned grids and incubated overnight, before infection with the EAV virus at a MOI of 10. Infected cells were incubated at 37°C for 12 hours prior to fixation. The culture medium was replaced with PBS containing 3.7% formaldehyde for 10 minutes at room temperature. The fixative was then removed, and cells were washed three times with PBS to remove residual formaldehyde. Following inactivation, grids were immediately blotted from the backside with filter paper and plunge-frozen in liquid ethane using a Leica GP2 plunge freezer, with 95% humidity and a blot force of 0 for 8 seconds. Vitrified grids were stored in liquid nitrogen until further use.

#### Cryo-focused ion beam milling of EAV-infected cells

Grids with EAV-infected Huh-7 cells were clipped into cryo-FIB AutoGrids™ (Thermo Fisher Scientific). Lamella preparation was performed on a dual-beam FIB/SEM Aquilos 2 microscope (Thermo Fisher Scientific Aquilos 2). The chamber and stage were maintained at below –170°C. To protect the biological material from ion beam damage, grids were sputter-coated with platinum for 30 s, followed by 30 s of gas injection system coating and additional 30s of platinum. For each target cell, lamella were milled by a gallium ion beam source at a 12° milling angle, at 30 kV in sequential steps: 1 nA for milling stress relief cuts and rough milling, 300 pA for medium milling, 100 pA for fine milling, and 30 pA for polishing to generate final thickness around 150-200 nm An additional sputtering coating was applied to lamella after final polishing for 5 s at 7 mA. Grids containing milled lamellae were stored in liquid nitrogen Dewars prior to tilt series collection. For EAV-infected cells, 49 lamellas were generated.

#### Cryo-ET data collection and tomogram reconstruction

Grids were loaded into a 300 kV Titan Krios G4 cryo-transmission electron microscope (Thermo Fisher Scientific) and equipped with a Selectris X energy filter. Tilt series were collected automatically using SerialEM software and PACEtomo script. Data were acquired at a nominal magnification of 53,000×, corresponding to a calibrated pixel size of 2.417 Å at the specimen level. Using a dose-symmetric tilt scheme starting at +12°, images were recorded from –50° to +64° with a 2° increment and 2 electron/Å^2^/tilt. The target defocus was set between –3 µm and –4 µm. Tilt series processing, including frame alignment, tilt series alignment, and 3D reconstruction, was performed using AreTomo v3.0 with the following key parameters: tilt axis rotation angle of –85°, tomogram alignment thickness of 1200 pixels, and a DarkTol threshold of 0.6. Motion correction and tilt series alignment were performed using a default patch-based method. The aligned tilt series were reconstructed into tomograms using weighted back-projection (Wbp 1), with a final binning factor of 4, resulting in a pixel size of 9.668 Å for initial visualization. Subsequent denoising and missing-wedge correction were performed on the AreTomo-reconstructed tomograms using IsoNet2^62^.

#### STA and segmentation analysis

1073 particles were manually picked from 22 denoised tomograms using the ArtiaX 0.6.0^49^ plugin in ChimeraX^50^, with their orientations predefined as perpendicular to the membrane. The particle poses were imported to RELION 4.0^47^ for further 3D classification and 3D refinement, applying C6 symmetry. These particles (bin4) were subjected to 3D classification and refinement in RELION 4.0. Particles from good class (306 particles) were refined with 2× binned pseudosubtomograms with C6 symmetry applied. Following further postprocessing, a final reconstruction of the pore complex was obtained at an overall resolution of 21 Å. Membrane segmentation was performed on the denoised and missing-wedge corrected tomogram using the v10_beta model of MemBrain-Seg^63^. RNA and RNP-like elements were manually annotated for 3∼5 slices of the tomogram in Ais 1.0.47^64^. These annotations were then cropped and used to train two separate neural networks in Ais, based on default VGGM architecture and training parameters, to generate prediction volumes. The membrane, RNA, and RNP-like elements prediction volumes were subsequently imported into ChimeraX for visualization and post-processing.

To improve visual clarity, the volumes were subjected to Gaussian filtering along the z-axis, followed by removal of small components using the “hide dust” function in ChimeraX. Ribosomes were picked by template matching against 80S ribosome map (EMD-44909) in emClarity 1.6.2^48^. Pore complexes were visualized using the final density map resolved by cryo-EM, with coordinates and orientations defined manually. Both ribosomes and pore complexes were low-pass filtered to 20 Å and imported into ChimeraX using ArtiaX 0.6.0^49^ for placement and visualization.

### Structural alignment and phylogenetic analysis

Protein structure searches were performed against Big Fantastic Virus Database (BFVD)^65^ on the Foldseek Seach Server^66^ (https://search.foldseek.com) in 3Di/AA and iterative search mode. The searches were conducted using the EAV nsp3 CTD, corresponding to residues 165–235, and the SARS-CoV-2 nsp4 CTD, corresponding to residues 410–493 of PDB entry 8YB7. Hits with Foldseek probability scores below 0.5 and entries that had been removed in UniProt were excluded. For strain with multiple hits, the hit with the highest probability score or lowest E-value was selected as the representative. After filtering, 33 hits were retained in addition to the two query structures. Multiple structural alignments used for downstream phylogenetic analysis are provided in **Extended Data Fig. 12**.

The aligned structural fragments were extracted according to the Foldseek alignment boundaries and used to generate a multiple structural alignment by FoldMason^67^ with default parameters. The amino-acid alignment and 3Di-state alignment were concatenated^68^ and used for phylogenetic inference in IQ-TREE 3.1.2^69^. The best-fit substitution models were selected using ModelFinder^70^. The selected models were Q.PFAM+G4 for the amino-acid partition and PMB+F+G4 for the 3Di-state partition. Branch support was assessed using 1000 ultrafast bootstrap replicates^71^ and 1000 SH-aLRT replicates. The unrooted tree was visualized using iTOL 7.5.1 (https://itol.embl.de/) and further refined in Adobe Illustrator.

### Data availability

Cryo-EM maps and corresponding atomic coordinates have been deposited in the Electron Microscopy Data Bank (EMDB) and the Protein Data Bank (PDB), respectively. Accession codes are as follows: EAV DMV pore complex by STA in C6 symmetry: EMDB EMD-XXXXX; EAV DMV pore complex by SPA in C3 symmetry: EMDB EMD-81134, PDB 27FZ. The raw images of isolated EAV DMV for cryo-EM SPA will be deposited to the EM Public Image Archive under accession code EMPIAR-XXXX. All other data supporting the findings of this study are available from the corresponding author on reasonable request.

## Acknowledgements

We thank Z.G. Yi at Fudan University (Shanghai, China) for the generous gift of the replicon vector which contain bicistronic reporter-selectable marker cassette. We also thank D.Y. Jin, Z.W. Ye and C.X. Wang for their advice in virus rescue and propagation; X. Fang for guidance in model building, data processing and structure analysis; and L.T. Fu for assistance with cryo-EM and cryo-ET data collection. The cryo-EM and cryo-ET data collection is accessed through Li Ka Shing cryo-EM laboratory under Centre for PanorOmic Sciences (CPOS) in The University of Hong Kong. The data processing is partially supported by the High-Performance Computing servers in CPOS in HKUMed. This work was supported by the National Natural Science Foundation of China – Young Scientists Fund (Category B, 32522006 to T.N.), Hong Kong Research Grant Council – Early Career Scheme (25129224 to X.Y.), Hong Kong Research Grant Council – General Research Fund (17122924 and 17127325 to T.N.), Research Grant Council – Young Collaborative Research Fund (C7065-25Y to T.N. and X.Y.) and HKUMed Research Collaboration Booster Fund as part of Tam Wing Fan Medical Development Fund.

## Author contributions

T.N. conceived and supervised the project and experiments. W.Z. isolated DMV and carried out cryo-EM SPA and cryo-ET STA of isolated DMVs with T.Y; W.Z., L.Z., Y.H., W.H. and T.N. analyzed cryo-EM/ET data; W.Z. and Y.H. built the atomic model; W.Z. and Y.G. prepared constructs and performed Western blots and co-IP; T.Y. constructed the EAV replicon for luciferase-based functional assays; T.Y. performed the virus rescue, propagation and inactivation validation of the infectious EAV virus; T.Y. and Y.H. performed the FIB-milling of EAV-infected cells and in situ cryo-ET analysis of EAV replication; Q.L. purified proteins; W.Z., W.H., and Y.W. analyzed the sequence and structure conservation; L.Z., Q.Y. and H.J. performed RT-SEM; W.Z., T.Y., X.Y. and T.N. drafted the initial manuscript and all authors contribute to manuscript.

## Main Figures

**Extended Data Table 1:** Cryo-EM data collection, refinement, and validation statistics.

|  | STA (EAV-infected) | STA (isolated recombinant) | SPA |
| --- | --- | --- | --- |
| **Data collection** |  |  |  |
| Microscope | TFS Krios | TFS Krios | TFS Krios |
| Voltage (kV) | 300 | 300 | 300 |
| Camera | Falcon 4i | Falcon 4i | Falcon 4i |
| Magnification | 53,000× | 81,000× | 130,000× |
| Pixel size (Å/pix) | 2.417 | 1.571 | 0.9557 |
| Total electron exposure/dose (e<sup>-</sup>/Å<sup>2</sup>) | 116* | 105, 99* | 40 |
| Defocus range (μm) | -3 to -4 | -1.5 to -4.5 | -0.5 to -1.5 |
| Energy filter slit width | 20 | 20 | 20 |
| Tilt-series (no.) | 73 (22)* | 1708 (1295)* | — |
| Tilt angle range (°) | -50 to 64** | -48 to 48** | — |
| Tilt step range (°) | 2 | 3 | — |
| Number of frames/image (EER) | 477 | 189/234/261 | 1197/1206 |
| Micrographs (no.) |  |  | 33,273 (21,185) |
| **Image processing** |  |  |  |
| Initial particles/subtomograms (no.) | 1073 | 385,650 | 2,149,570 |
| Final particles/subtomograms (no.) | 306 | 19,535 | 76,112 |
| Symmetry imposed | C6 | C6 | C3 |
| Map resolution (Å) | 21 | 7.5 | 3 |
| FSC threshold | 0.143 | 0.143 | 0.143 |
| Map resolution range (Å) | — | 6.5 to 8.5 | 2.5 to 5.0 |
| **Refinement** |  |  |  |
| Initial model used | — | — | N/A |
| Model resolution (Å) | — | — | 3.34 |
| FSC threshold (model) | — | — | 0.5 |
| Map sharpening B factor (Å<sup>2</sup>) | — | — | -72 |
| Model composition |  |  |  |
| Non-hydrogen atoms | — | — | 15230 |
| Protein residues | — | — | 2026 |
| Lipids | — | — | 0 |
| Ions | — | — | 4 (Zn) |
| B factor (Å<sup>2</sup>) |  |  |  |
| Protein | — | — | 34.24 |
| Ligands | — | — | 56.71 |
| R.m.s. deviations |  |  |  |
| Bond lengths (Å) | — | — | 0.007 |
| Bond angles (°) | — | — | 0.744 |
| Validation |  |  |  |
| MolProbity score | — | — | 1.37 |
| Clash score | — | — | 4.67 |
| Poor rotamers (%) | — | — | 1.15 |
| Ramachandran plot |  |  |  |
| Favored (%) | — | — | 97.71 |
| Allowed (%) | — | — | 2.39 |
| Disallowed (%) | — | — | 0 |

**Extended Data Fig. 1.**
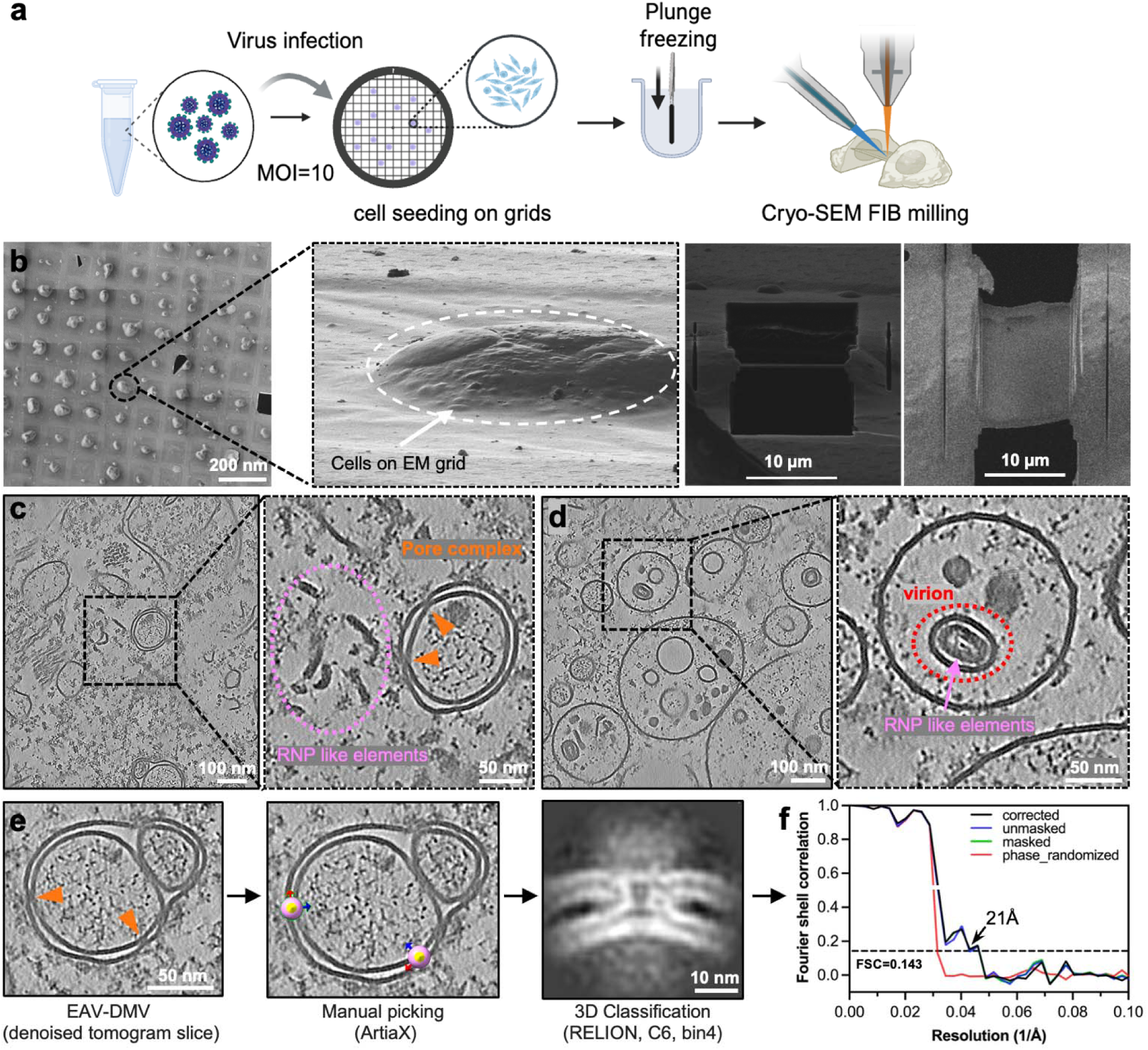
| Experimental pipeline for cryo-FIB milling and electron c yo-tomography of EAV-infected Huh7 cells. **a**, Schematic overview of the experimental workflow, including infection of Huh7 cells grown on EM grids, plunge freezing, and cryo-FIB milling. **b,** Representative cryo-SEM images showing Huh7 cells on EM grids and the preparation of a thin lamella for cryo-ET analysis. Dashed outlines indicate the selected cell region and the milled lamella. **c,** A representative tomogram slice of an EAV-infected cell showing DMVs; the enlarged view highlights a DMV pore complex (orange arrowheads) and surrounding RNP-like elements. **d,** Tomographic slice showing virion particles in infected cells; the enlarged view indicates a virion particle and inner RNP-like elements. **e**, Subtomogram averaging pipeline of DMV pore complex in situ. The pore particles were manually identified from denoised tomograms and their orientation of z axis is predefined as perpendicular to membrane. Extracted particles were then subjected to 3D classification in RELION with imposed C6 symmetry (bin4), yielding the representative pore density shown. **f**, Gold-standard half-map Fourier shell correlation curves for the final reconstruction. Corrected, unmasked, masked and phase-randomized FSC curves are shown. The estimated global resolution is 21 Å at the FSC = 0.143 criterion.

**Extended Data Fig. 2.**
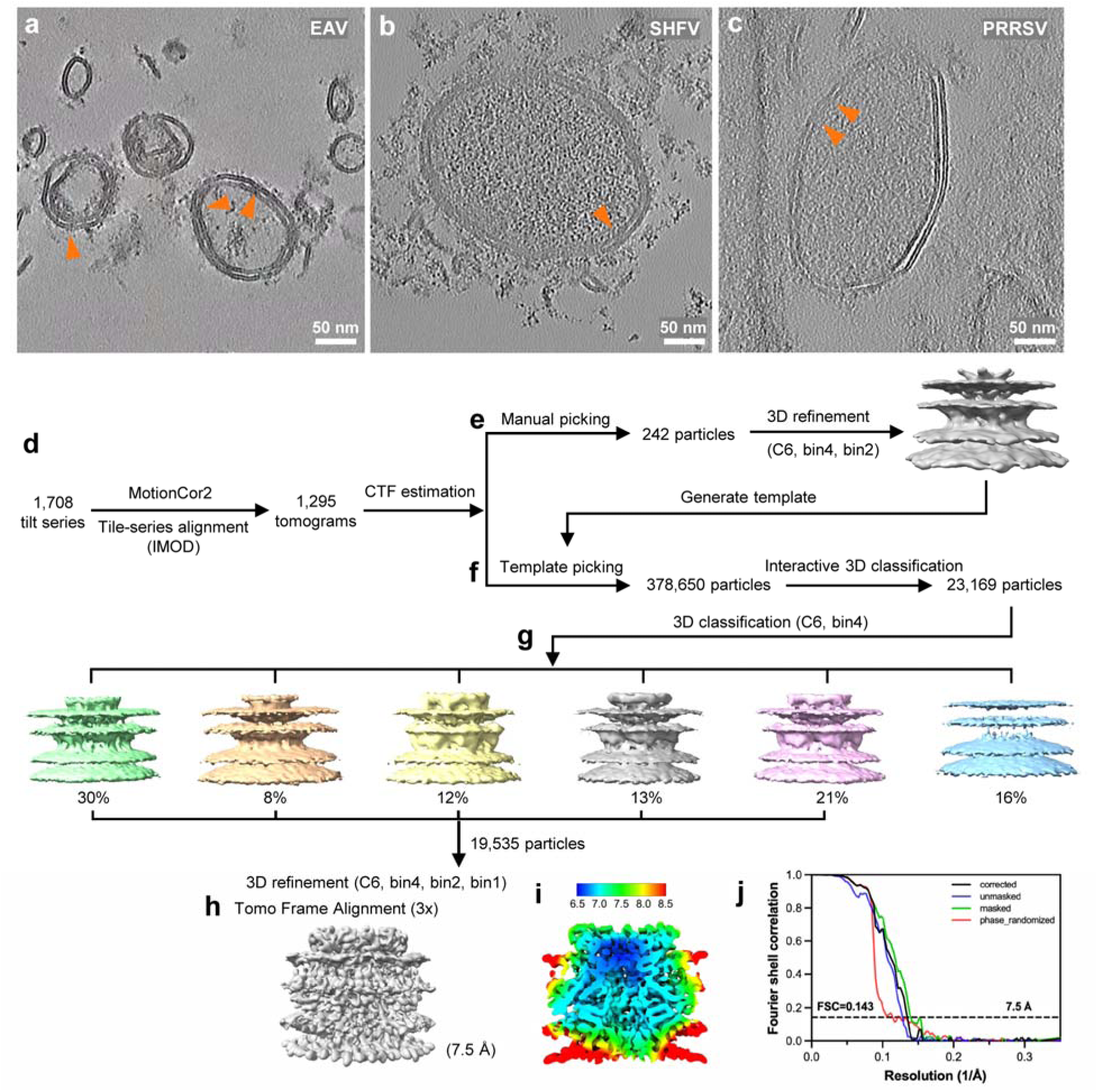
| Cryo-ET and subtomogram averaging of the EAV DMV pore complex. **a–c**, Representative tomographic slices showing purified arterivirus DMVs: EAV (**a**), SHFV (**b**) and PRRSV (**c**). **d,** Processing pipeline for STA of the isolated EAV nsp2–nsp3 DMV pore complex. 1,708 tilt series were motion-corrected, aligned and manual inspected, yielding 1,295 tomograms containing DMVs for further analysis. **e,** After CTF estimation in emClarity, an initial set of manually picked particles was used to generate a template, which was then applied for template matching. **f,** Template picking is performed in emClarity, which were subjected to interactive 3D classification in RELION for cleaning, resulting in 23,169 particles. **g,** Six representative 3D classes are shown with their relative particle distributions. **h,** Particles from the selected five classes were pooled, giving 19,535 particles for final 3D refinement with C6 symmetry using progressively bin4, bin2 and bin1 with three times of tomo-frame alignment. The final subtomogram average of the EAV DMV pore complex reached an overall resolution of 7.5 Å. **l,** Local-resolution map of the final reconstruction from side view cross-section, coloured according to the indicated resolution scale. **j,** Gold-standard half-map Fourier shell correlation curves for the final reconstruction. Corrected, unmasked, masked and phase-randomized FSC curves are shown. The estimated global resolution is 7.5 Å at the FSC = 0.143 criterion.

**Extended Data Fig. 3.**
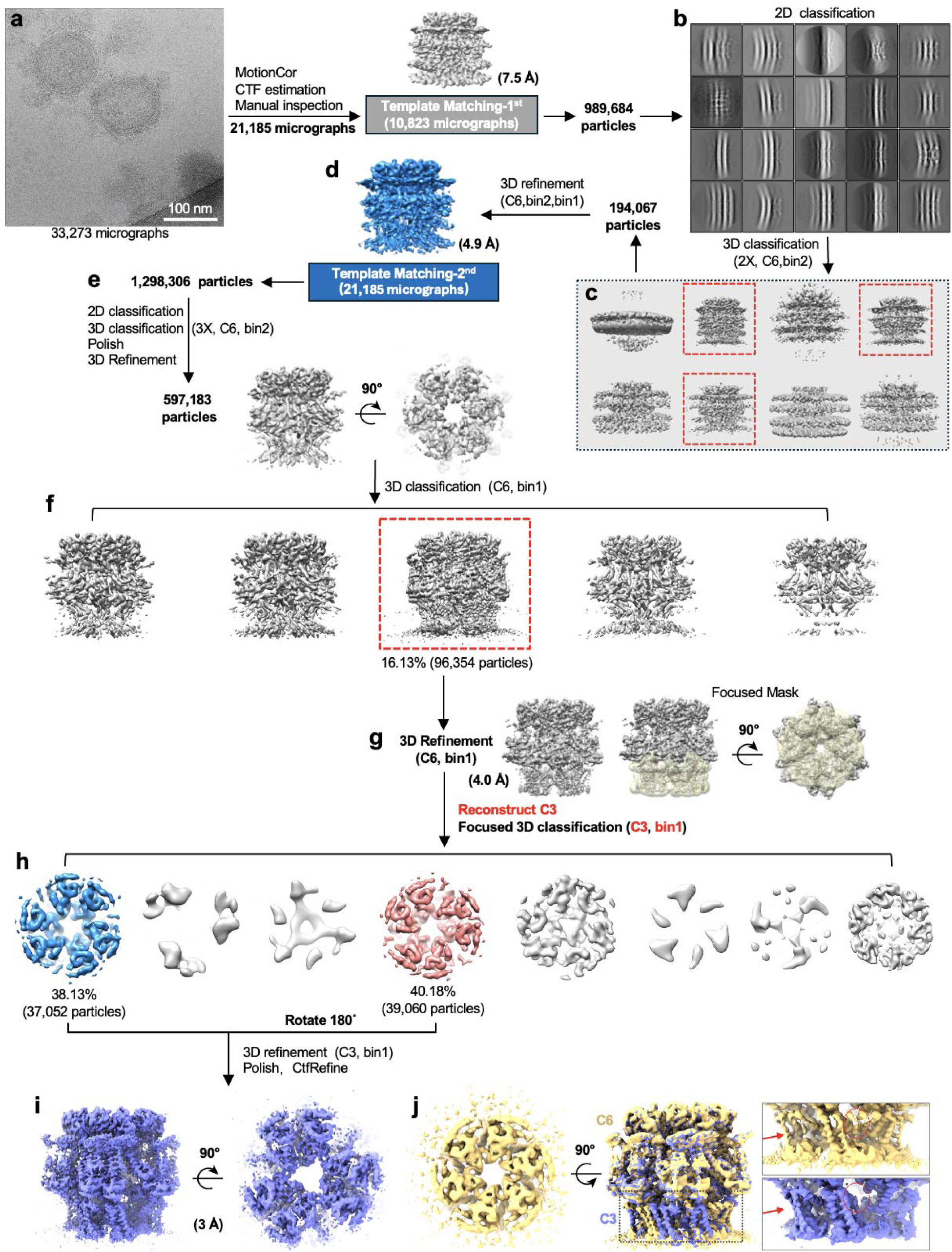
| Cryo-EM and single-particle analysis of the isolated EAV DMV pore complex. **a**, A representative cryo-EM micrograph with two isolated EAV DMVs. **b-d**, Workflow for the first round of template matching and classification. A total of 33,273 cryo-EM movies were collected, motion-corrected and subjected to CTF estimation, followed by manual inspection to retain 21,185 micrographs containing DMVs. The 7.5 Å STA map was used as an initial template for template matching on a subset of 10,823 micrographs, yielding 989,684 particles. **b-c**, Representative 2D classification skipping alignment, whose particles from potential classes were subjected to two rounds of 3D classification with C6 symmetry. Classes corresponding to well-defined DMV pore complexes (highlighted with red dashed boxes) were selected for further 3D refinement with C6 symmetry at bin2 and bin1 subsequently, yielding a 4.9 Å reconstruction as **d** as an improved template for a second round of template matching. **e**, Workflow of the second-round template matching and processing. A total of 1,298,306 particles at bin2 were subjected to one round of 2D classification, three rounds of 3D classification with C6 symmetry, Bayesian polishing and 3D refinement. A subset of 597,183 particles with well-defined structural features was selected and extracted at bin1 for 3D refinement, with representative side and top views of the refined map. **f**, One class with the clearest secondary-structure features was selected from 3D classification (bin1, C6 symmetry) for subsequent refinement (red dashed box). **g**, 3D refinement performed on selected class at C6-symmetry followed by focused 3D classification with a local mask covering the nsp3 region at C3 symmetry. **h**, Focused 3D classification showing two major classes with clear nsp3 CTD density were selected. One class was rotated by 180° and combined with the other for final refinement. **i**, Final C3-symmetric reconstruction after 3D refinement, Bayesian polishing and CTF refinement, reaching an overall resolution of approximately 3.0 Å. Side and top views are shown. **j**, Comparison of the C6-symmetric reconstruction and the C3-focused reconstruction.

**Extended Data Fig. 4.**
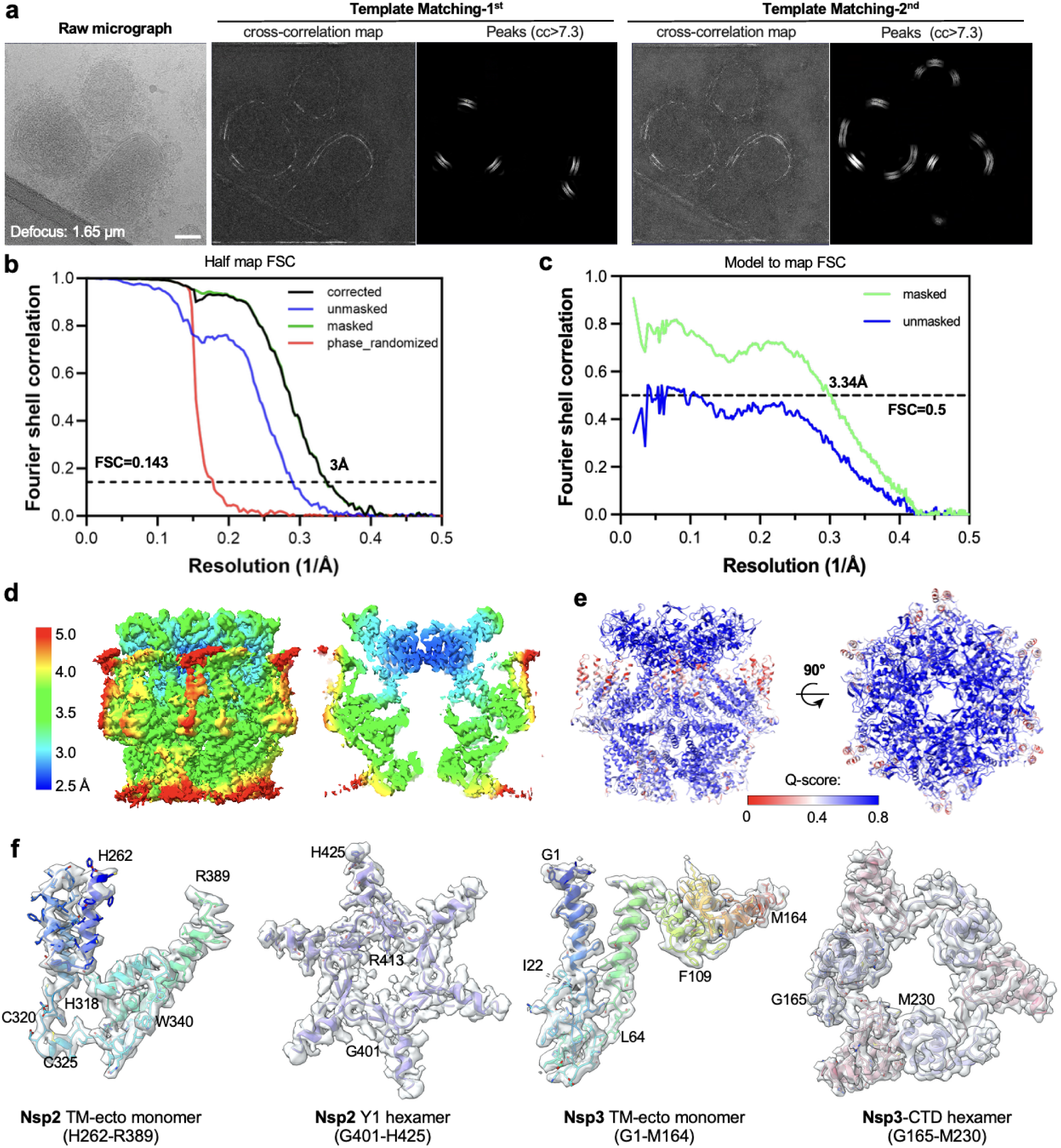
| Template matching, FSC analysis and local-resolution estimation. **a**, Representative cryo-EM micrograph and its corresponding template-matching results. Micrograph defocus: 1.65 µm. Cross-correlation maps and detected peaks from the first and second rounds of template matching are shown. Peaks were shown using a cross-correlation threshold of >7.3. The second round of template matching identified additional particles at the same threshold. **b,** Gold-standard half-map Fourier shell correlation curves for the final reconstruction. Corrected, unmasked, masked and phase-randomized FSC curves are shown. The estimated global resolution is 3.0 Å at the FSC = 0.143 criterion. c, Model-to-map FSC curves calculated using masked and unmasked maps. The masked model-to-map FSC indicates an estimated resolution of 3.34 Å at the FSC = 0.5 criterion. **d**, Local-resolution estimation of the final reconstruction, shown in surface representation from side view, cross-section and top views. **e**, Atomic model coloured by local Q-score values calculated in ChimeraX and shown in two orthogonal orientations from 0–0.8 (red to blue). **f,** Representative transparent cryo-EM density map and fitted atomic models coloured in ribbons for selected regions of the complex.

**Extended Data Fig. 5.**
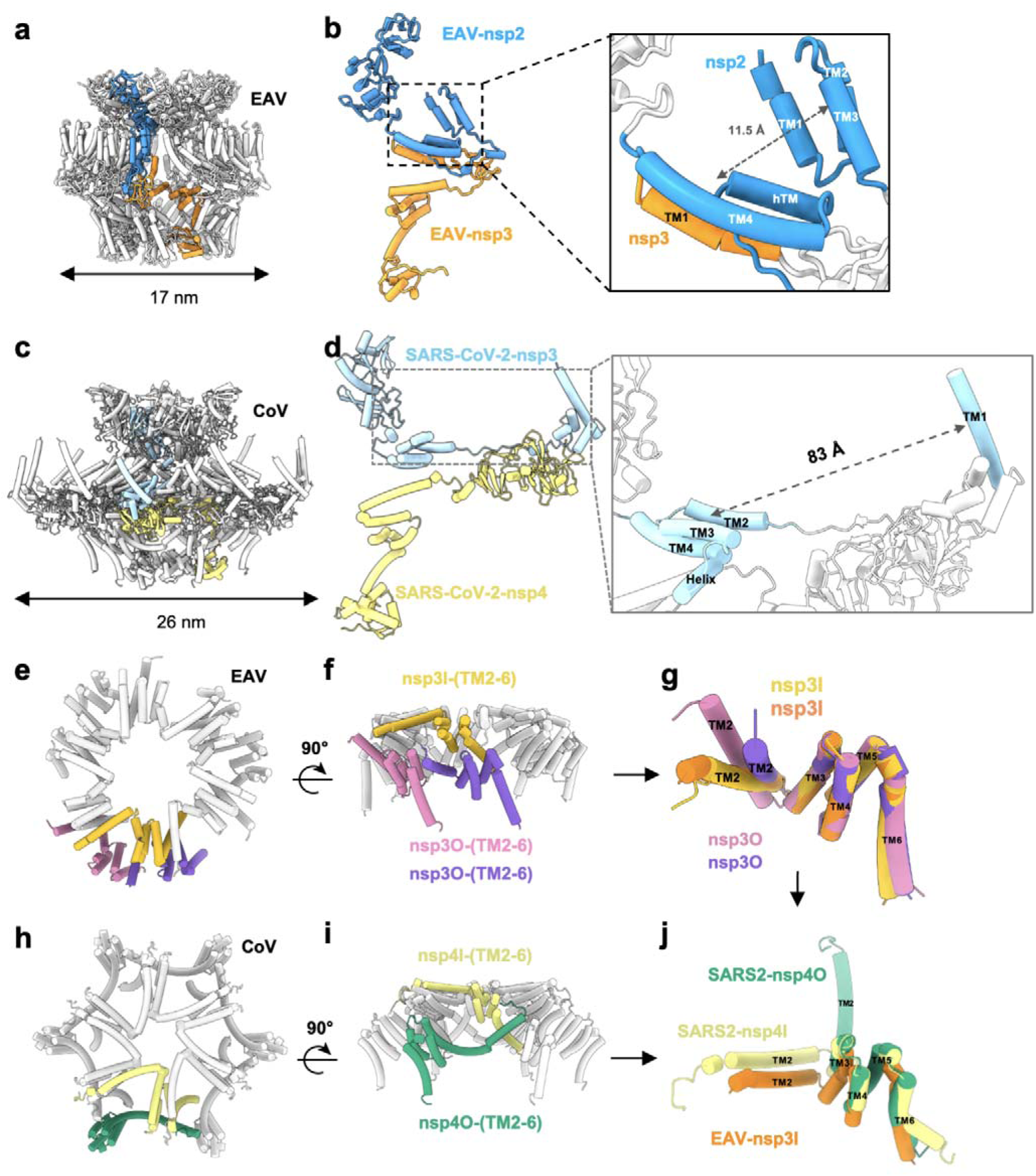
| Comparison of ectodomains and transmembrane helices from arterivirus and CoV. **a,c**, Overall architecture of the EAV nsp2–nsp3 complex in **a** or SARS-CoV-2 nsp3-nsp4 complex (PDB:8YB7) in **c**, highlighted with one EAV nsp2(blue)-nsp3(orange)/ CoV nsp3(cyan)-nsp4(yellow) heterodimeric unit. **b,d,** Representative extracted views of the EAV nsp2–nsp3 unit from **a** shown in **b**, or CoV nsp3-nsp4 unit from **c** shown in **d**, with an enlarged view of the transmembrane region in the right panel. The inset highlights the relative arrangement of transmembrane helices in DMV outer membranes from arterivirus and CoV, with approximate inter-helical distances are indicated. The distances are measured in ChimeraX by defining the center of gathered helices. **e, f,** Top (**e**) and side (**f**) view of the transmembrane-helix arrangement from arterivirus nsp3 within the oligomeric assembly. Coloured helices indicate selected one representative TM2–TM6 segment, while the remaining helices are shown in grey. **g,** Superposition of selected TM2–TM6 bundles of arterivirus nsp3 from **f**, including nsp3I and nsp3O, highlighting the conserved arrangement of TM3-TM6 and different directions of TM2. **h, i,** Views of the corresponding transmembrane-helix arrangement from CoV nsp4 within the oligomeric assembly, with one representative inner nsp4 (nsp4I, light yellow) and outer nsp4 (nsp4O, green) Panel **h** shows a top view and panel **i** shows a side view after a 90° rotation. **j,** Structural comparison of TM2–TM6 bundles from SARS-CoV-2 nsp4 (nsp4I, light yellow; nsp4O, green) and one representative EAV nsp3 (nsp3I, orange).

**Extended Data Fig. 6.**
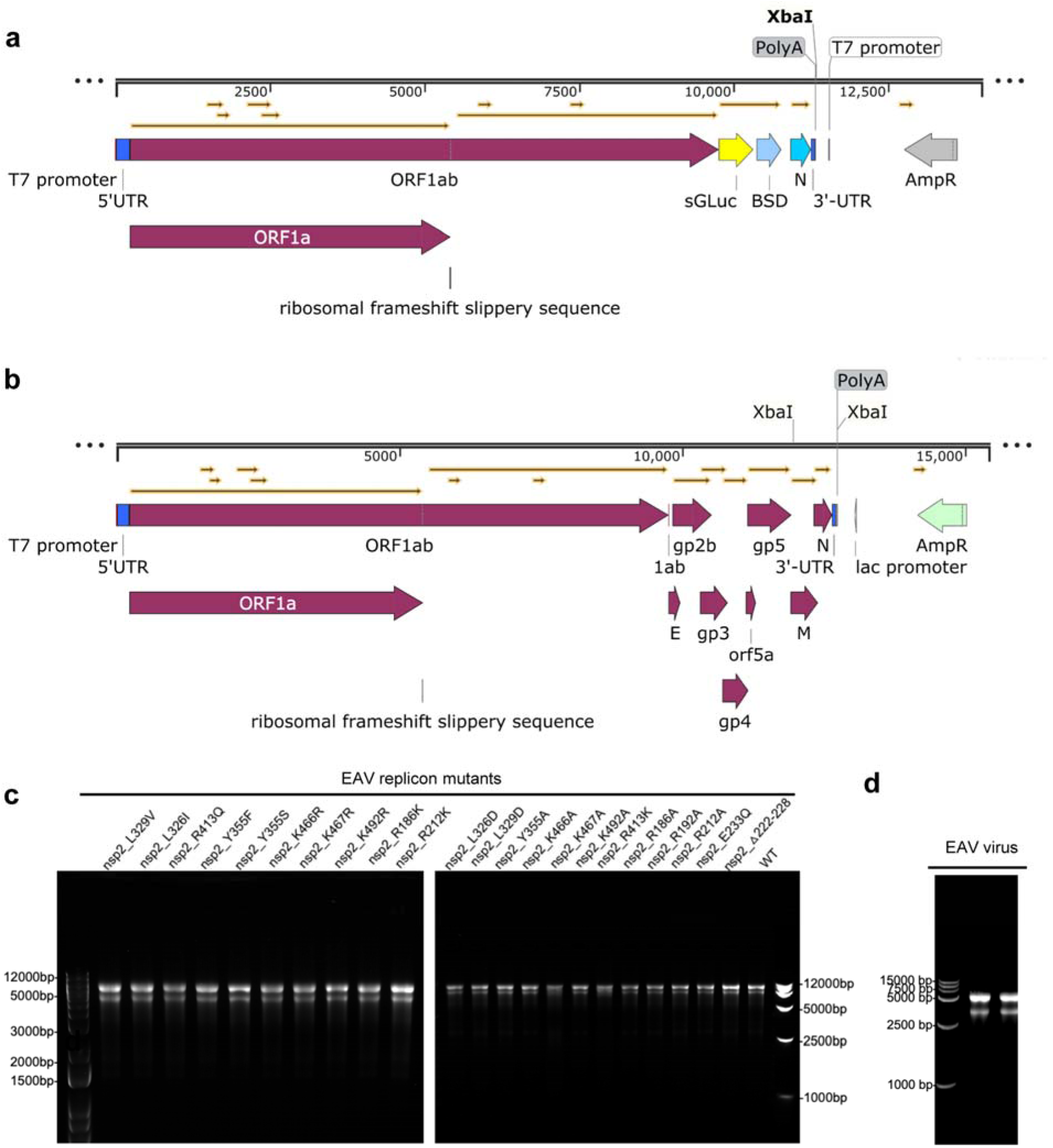
| Schematic of construction of the EAV replicon and full-length EAV clone, in vitro transcription, and RT–PCR verification. **a.** Genome organization of the virulent Bucyrus strain of EAV are shown at the top. A T7 promoter was placed upstream of the 5′ UTR, and a poly(A)20 tract plus XbaI site was introduced at the 3′ end for IVT and plasmid linearization. An expression cassette containing secreted *Gaussia* luciferase (sGluc), foot-and-mouth disease virus (FMDV) 2A peptide and blasticidin (BSD) was added to replace the structural proteins except N. **b,** Schematic representation of the full-length EAV cDNA clone. Similar to the replicon shown in a, a T7 promoter was inserted upstream of the 5′ UTR and a poly(A) tract plus an XbaI site was introduced at the 3′ end for plasmid linearization and in vitro transcription. The full set of structural protein genes is retained. **c.** Agarose gel analysis of the indicated mutant replicon constructs and WT control. **d,** Agarose gel analysis of the full-length EAV transcription product.

**Extended Data Fig. 7.**
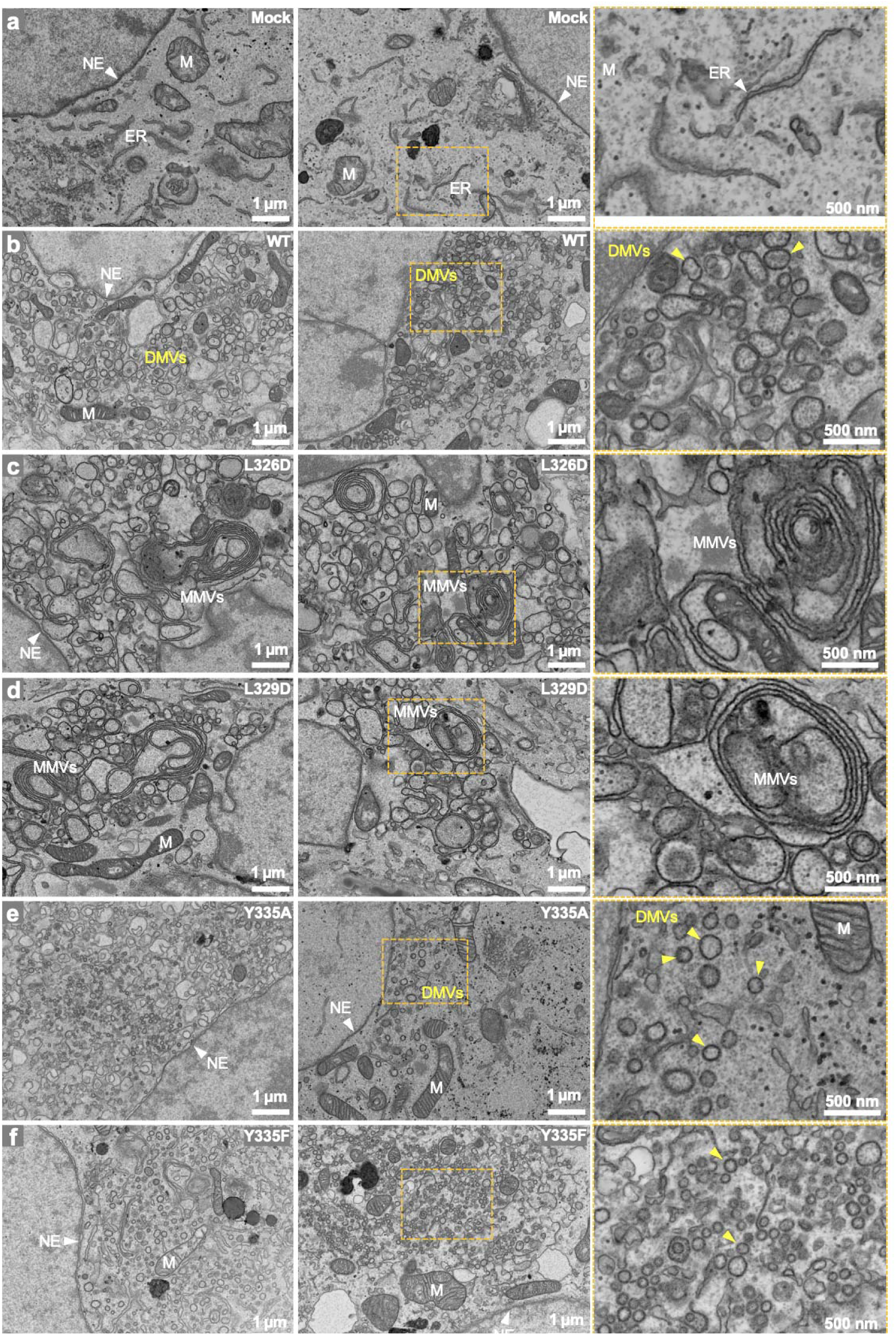
| SEM images of nsp2-nsp3 WT and nsp2 interface mutant in transfected COS-7 cells. **a.** Representative SEM images of untransfected COS-7 cells, displaying normal endoplasmic reticulum (ER) organization. **b.** SEM images of COS-7 cells at 36-h post transfection with WT nsp2-nsp3 construct, exhibiting the formation of DMVs compared to the control in (**a**). **c-f**. Representative images of COS-7 cells transfected with nsp2-nsp3 constructs containing nsp2 interface mutations: L326D (**c**), L329D (**d**), Y335A (**e**), Y335F (**f**). Two representative images are shown for each experiment group with enlarged views (orange boxes) on the right panel. NE, nuclear envelope (white arrowheads); ER, endoplasmic reticulum; M, mitochondria; DMVs, double-membrane vesicles (yellow arrowheads); MMVs: multi-membrane vesicles. Scale bars are as indicated in the images.

**Extended Data Fig. 8.**
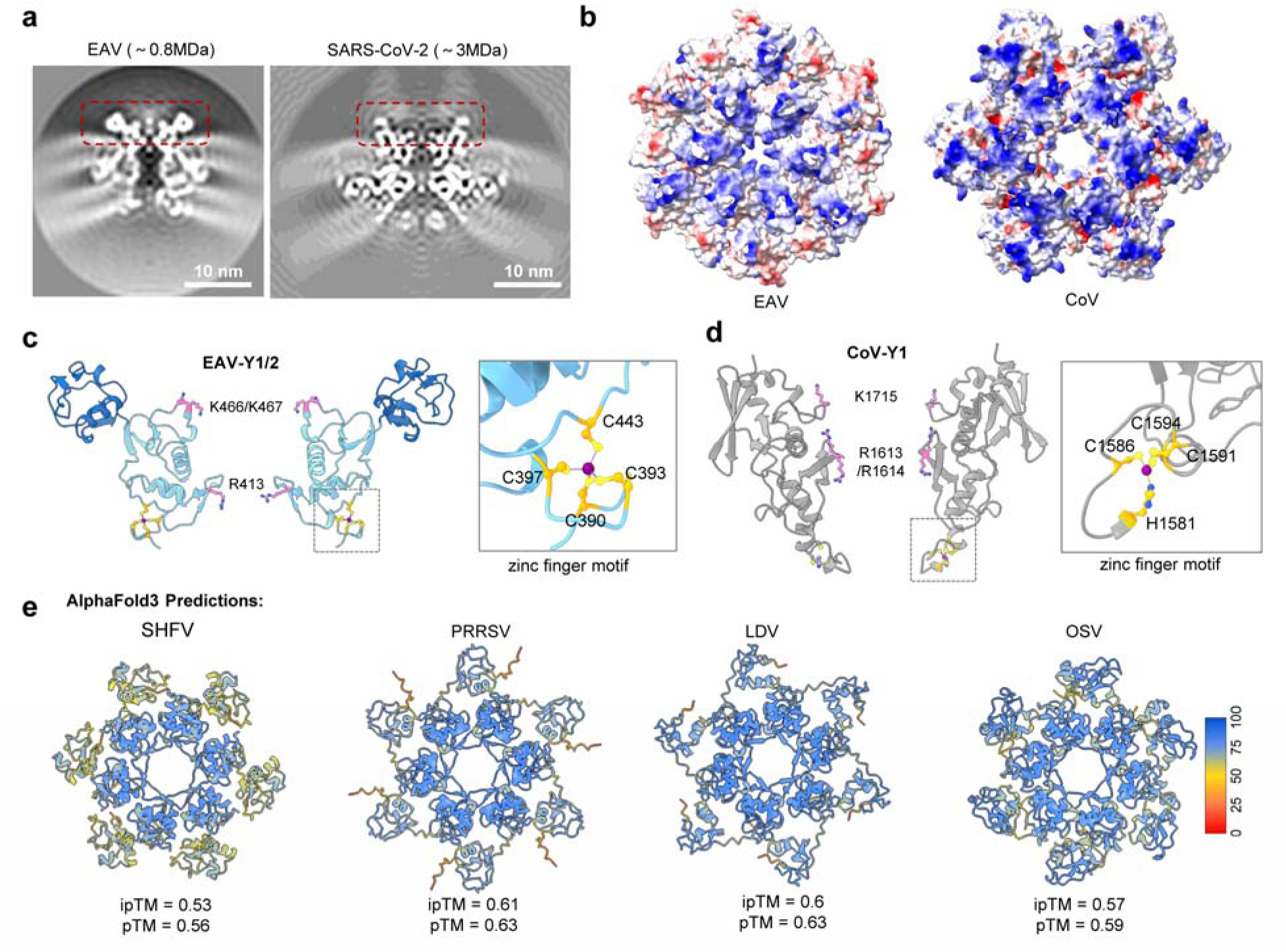
| Nsp2-Y1 domain conservation and hexamer formation. **a**, Cross-sections of the EAV DMV pore complex and the SARS-CoV-2 nsp3-nsp4 mini-pore complex (EMD-39113, both lowpass filtered to 12 Å). Approximate molecular masses are indicated, and red dashed boxes mark the Y1-hexamer. **b,** Bottom views of the EAV and CoV DMV pore Y1 oligomers coloured by electrostatic charge in ChimeraX. Blue to red: positive to negatively charged. **c,** EAV nsp2 Y1/Y2 shown in two orientations, with Y1 in cyan and Y2 in blue. Positively charged residues lining the central pore are highlighted in pink. The inset shows the zinc-finger motif, with conserved cysteines in yellow and the zinc ion as a magenta sphere. **d,** Coronavirus Y1-domain model shown in two orientations, with the conserved zinc-finger motif enlarged in the inset. **e,** AlphaFold3 predictions of hexameric nsp2 C-terminal domain assemblies from SHFV, PRRSV, LDV and OSV. Models are coloured by predicted confidence score, with ipTM and pTM values indicated below.

**Extended Data Fig. 9.**
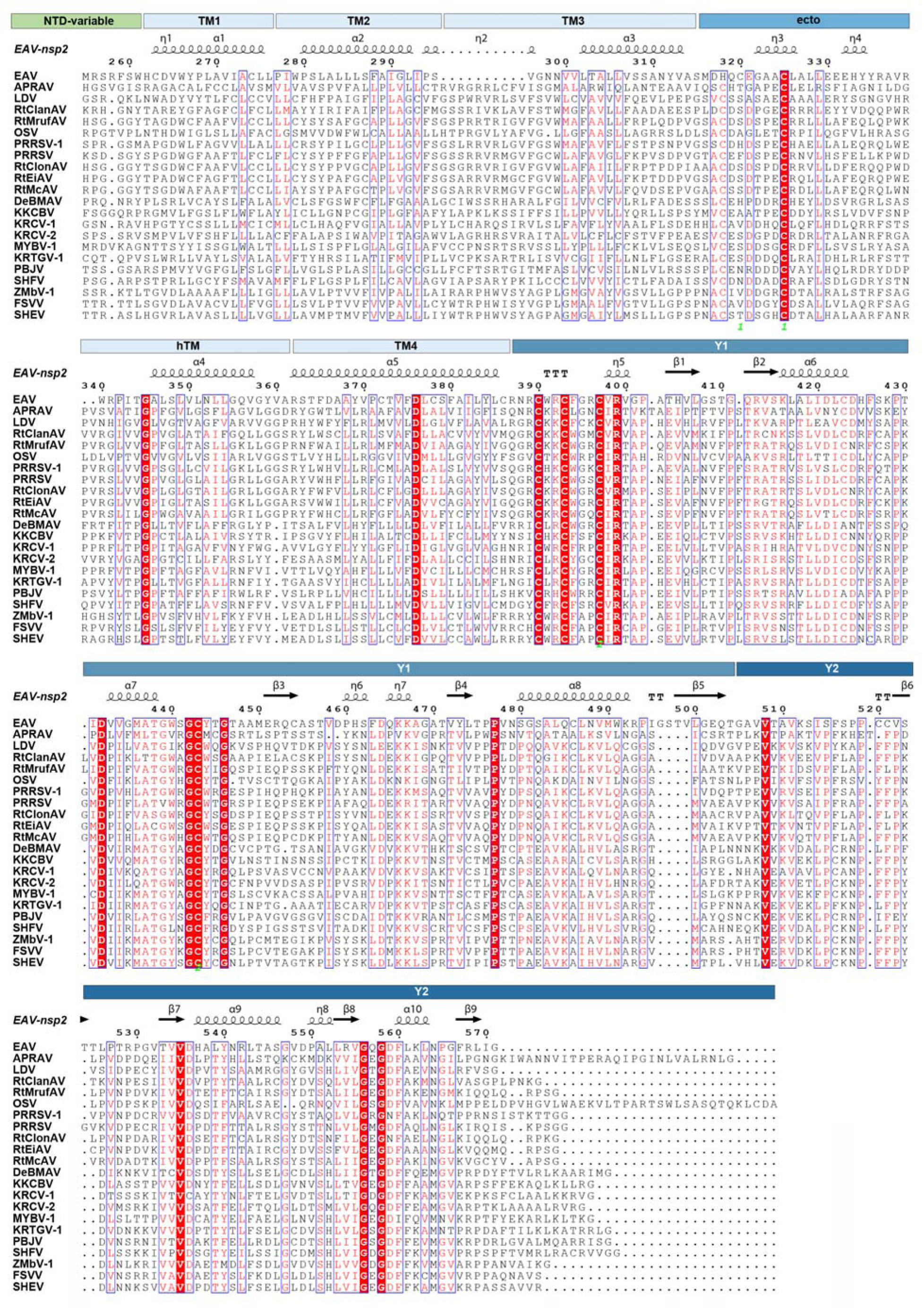
| Sequence alignment of arterivirus nsp2. Multiple sequence alignment of representative arterivirus nsp2 proteins, annotated with TM1–TM4, ectodomain, hTM, and Y1/Y2 regions, and the NTD-variable region are not shown here. Conserved and similar residues are highlighted in the alignment. Sequences are labelled with virus names with their Uniprot number shown here: OSV-1, A0A1Z2RX77; EAV, P19811; APRAV, A0A0B5JQL5; SHFV, Q68772; SHEV, A0A0F6PT34; FSVV, A0A161D9J9; ZMbV-1, A0A167L705; KRCV-2, X2D5J6; DeBMAV, A0A0B6C113; KRTGV-1, L0CQK2; PBJV, A0A0G2UIN6; KCCBV, A0A0Y0BQ57; MYBV-1, A0A089H3C1; KRCV-1, X2D6S1; PRRSV-2, AY150312.1; RtMruFAV, A0A0U2M952; RtClonAV, A0A1L6Z3P9; RtEiAV, A0A2H4MWP6; PRRSV-1, Q04561; RtMcAV, A0A2H4MWN4; LDV, Q83017; and RtClanAV, A0A2H4MXL2.

**Extended Data Fig. 10.**
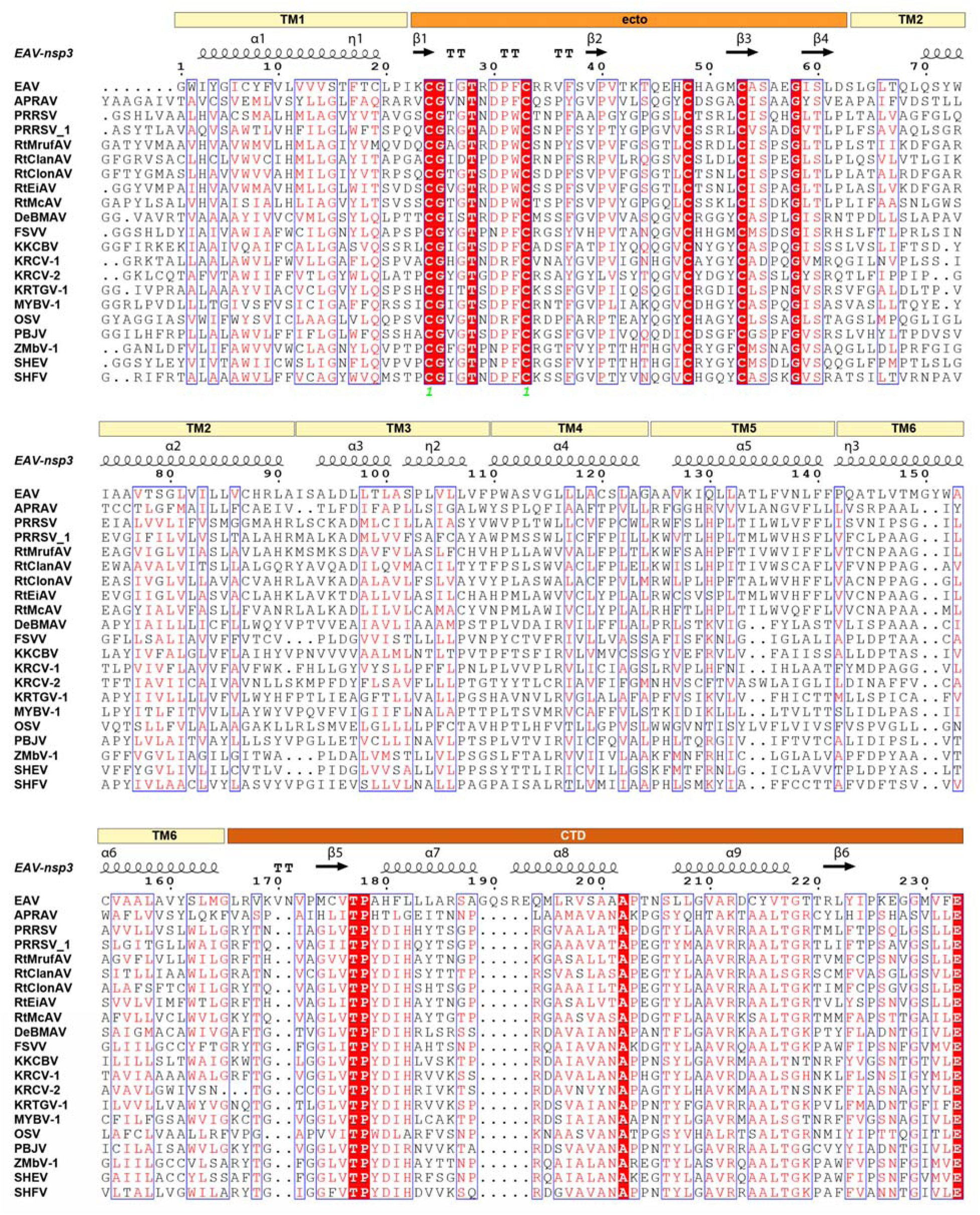
| Sequence alignment of arterivirus nsp3. Conserved and similar residues are highlighted in the alignment. Sequences are labelled with virus names with their Uniprot number shown here: OSV-1, A0A1Z2RX77; EAV, P19811; APRAV, A0A0B5JQL5; SHFV, Q68772; SHEV, A0A0F6PT34; FSVV, A0A161D9J9; ZMbV-1, A0A167L705; KRCV-2, X2D5J6; DeBMAV, A0A0B6C113; KRTGV-1, L0CQK2; PBJV, A0A0G2UIN6; KCCBV, A0A0Y0BQ57; MYBV-1, A0A089H3C1; KRCV-1, X2D6S1; PRRSV-2, AY150312.1; RtMruFAV, A0A0U2M952; RtClonAV, A0A1L6Z3P9; RtEiAV, A0A2H4MWP6; PRRSV-1, Q04561; RtMcAV, A0A2H4MWN4; LDV, Q83017; and RtClanAV, A0A2H4MXL2.

**Extended Data Fig. 11.**
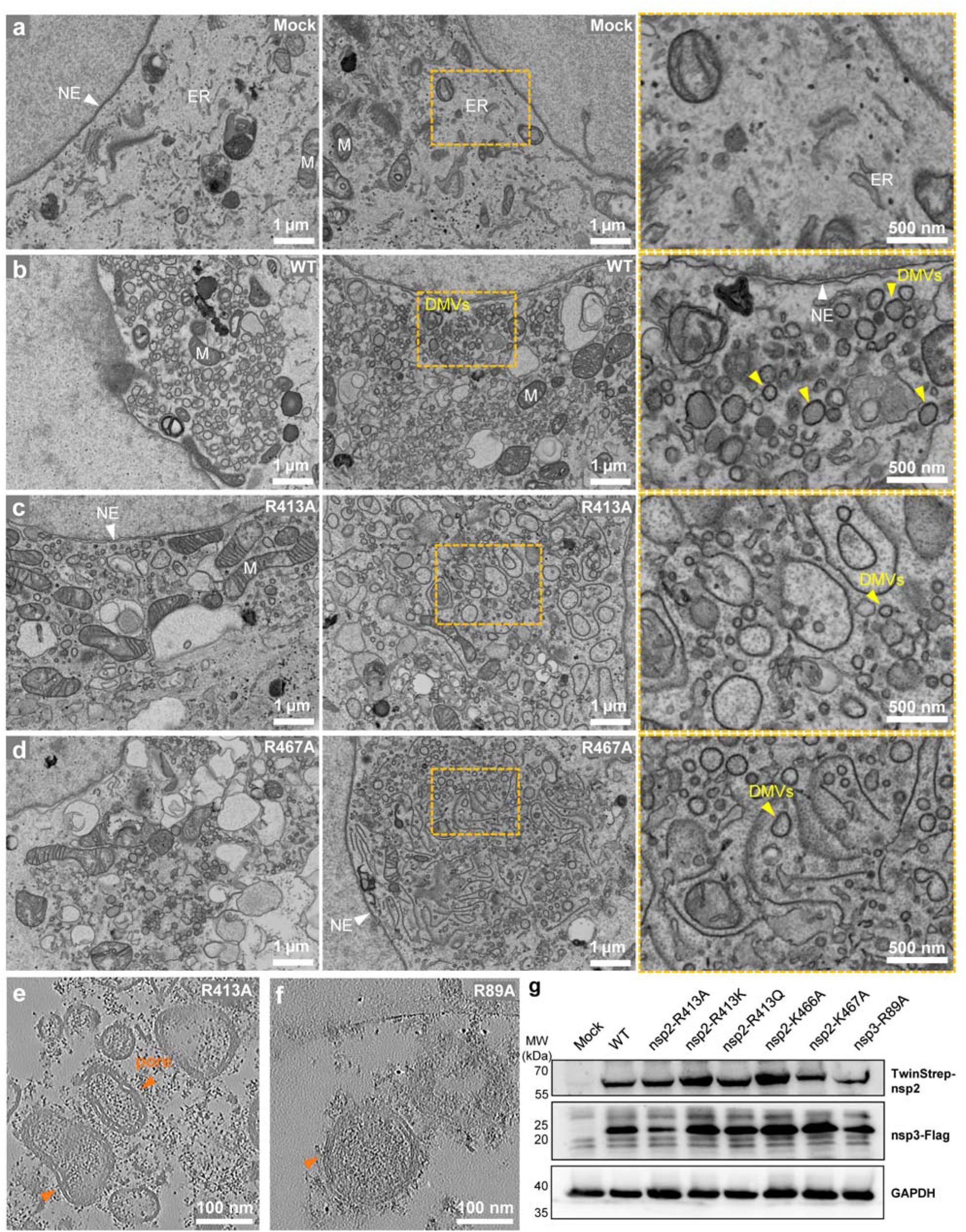
| Effects of nsp2 pore residue mutations on DMV and pore formation. **a-d**, Representative SEM images of COS-7 cells transfected with wild-type (WT) nsp2-nsp3 (**b**), nsp2 R413A (**c**) and nsp2 R467A (**d**) compared to that of untransfected cells in (**a**) at 36 h post-transfection timepoint. Two representative images are provided for each experimental condition with enlarged views (orange boxes) on the right panels. Abbreviations are as follows: NE, nuclear envelope (white arrowheads); ER, endoplasmic reticulum; M, mitochondria; DMVs, double-membrane vesicles (yellow arrowheads). Scale bars are as indicated in the images. **e-f**, Representative tomographic slices of purified EAV mutant DMVs from transfected HEK293F cells. **g**, Western-blot analysis of lysates from COS-7 cells that transfected with WT or nsp2 mutant plasmids, showing the expression of both EAV nsp2 and nsp3 proteins.

**Extended Data Fig. 12.**
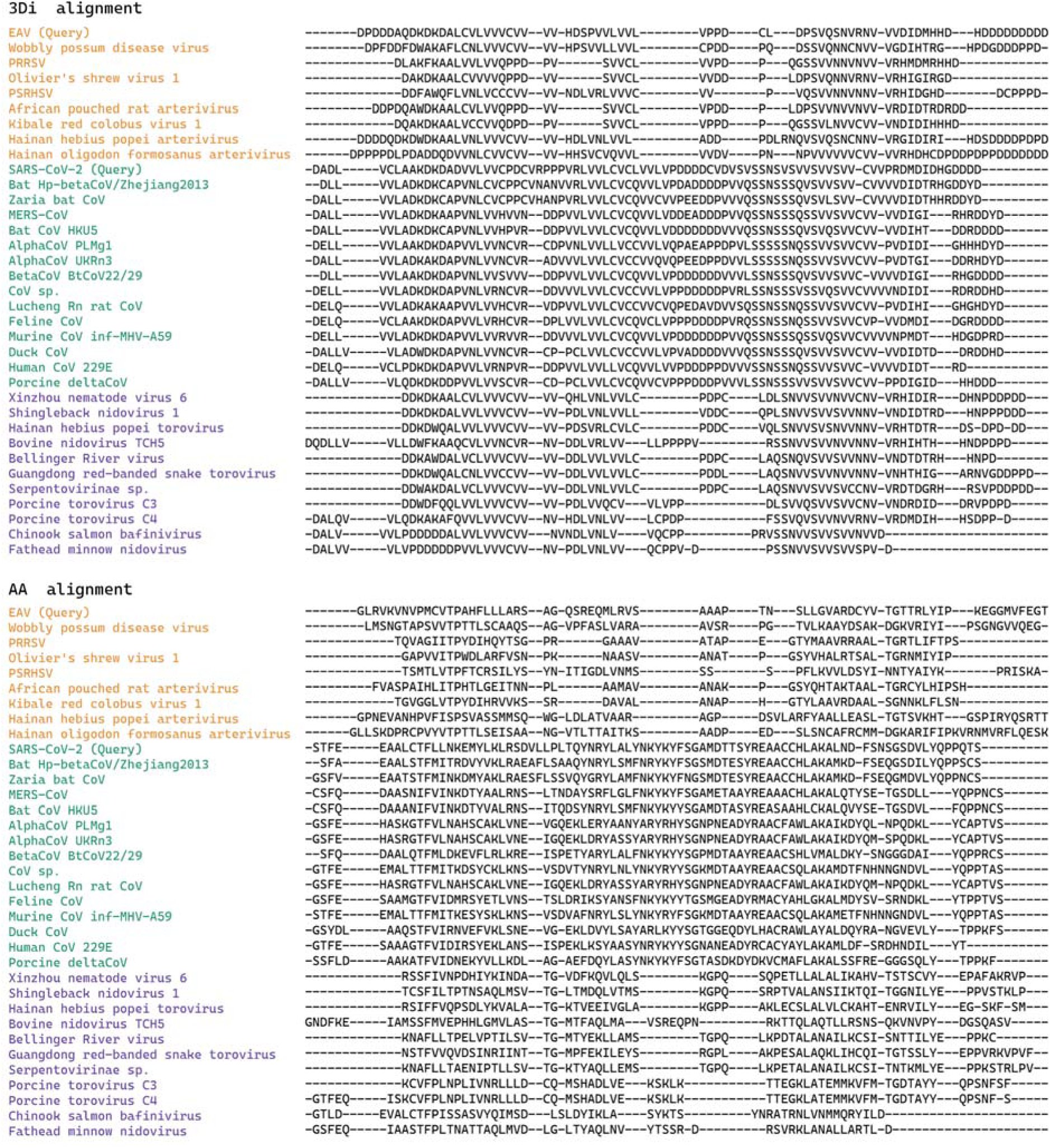
| Multiple structure alignment of EAV and SARS-CoV-2 CTD domain. Alignment was done by Foldmason using “--match-ratio 0.9 –-filter-msa 1 –-gap-open aa:25,nucl:25 –-gap-extend aa:2,nucl:2 –-report-paths 0 –-report-mode 2”. msaLDDT is 0.613063. Curated hit tables is provided as Supplementary Table 1. Abbreviation: PRRSV – Porcine reproductive and respiratory syndrome virus; PSRHSV – Pelodiscus sinensis respiratory and hemorrhagic syndrome virus; MERS-CoV – Middle East respiratory syndrome-related coronavirus.

